# Rapid Remodeling of Human White Adipose Tissue Following Bariatric Surgery

**DOI:** 10.64898/2026.04.09.717542

**Authors:** Zinger Yang Loureiro, Gregory P. Westcott, Anton Gulko, Adam Essene, Zhibo Zhou, Wenle Liang, Christopher Jacobs, Soumya Nagesh, Nadejda Bozadjieva-Kramer, Randy J. Seeley, William Gourash, Joseph J. Loureiro, Lori L. Jennings, Robert E. Gerszten, Anita Courcoulas, Linus T. Tsai, Margo P. Emont, Evan D. Rosen

## Abstract

Bariatric surgery induces profound weight loss and improvement of obesity-associated metabolic dysfunction. Recent studies have shown that adipose tissue undergoes remodeling after weight loss, characterized by a reduction in proinflammatory immune cells, increased vascularization, and a shift in the adipocyte transcriptome, but these studies focused on time points long after surgery. We performed single nucleus RNA-seq (snRNA-seq) in subcutaneous white adipose tissue (SAT) samples from subjects with obesity undergoing bariatric surgery, collected at baseline and at one, six, and twelve months after surgery. We identify profound remodeling of SAT within the first month after surgery, characterized by a surge in lipid-associated macrophages and sharp reductions in specific populations of adipocytes, adipose stromal and progenitor cells (ASPCs), and endothelial cells. Transcriptional profiles strongly suggest that some adipocytes undergo apoptosis soon after surgery, while new adipocytes are generated by *de novo* differentiation. Mechanistically, the data are consistent with a model whereby coordinated early loss of a hedgehog signaling axis between endothelial cells and an anti-adipogenic population of ASPCs known as adipose regulatory cells (Aregs; ASPC^PTCH2^) enables a transient burst of adipogenesis to occur. Interestingly, very few of these early features seen in human subjects are represented in a mouse model of surgical weight loss.

## INTRODUCTION

Obesity, one of the most common medical conditions of the 21st century, is associated with a wide array of adverse health consequences, including type 2 diabetes (T2D), cardiovascular disease, and certain cancers, among others. Traditional approaches like dietary control and exercise have, in general, been unsuccessful in treating excess body weight. Until the recent development of drugs based on glucagon-like peptide 1 receptor (GLP1R) agonism, bariatric surgery was the only effective therapy for obesity. In fact, the weight loss-promoting effects of bariatric surgery may be longer lasting and more cost effective than even current forms of pharmacotherapy, and many patients still opt for bariatric surgery as a preferred option for weight loss.^1^

Adipose tissue comprises multiple cell types, including the major parenchymal cell type, the adipocyte. In addition, adipose tissue contains large numbers of adipose stromal and progenitor cells (ASPCs), vascular cells (blood and lymphatic endothelial cells, pericytes, and vascular smooth muscle cells), and both myeloid and lymphoid immune cells.^2,3^ In addition, each of these major cell populations encompass several subpopulations, which differ in their gene expression patterns, and presumably in their function as well. During the development of obesity, the composition of adipose tissue is altered, most notably reflecting the influx of proinflammatory immune cells.^4^ In concert with these compositional changes, the metabolic activity of adipose tissue becomes deranged in the obese state. Studies in rodents and humans have pointed to several drivers of adipose dysfunction, including inflammation, activation of genes associated with senescence, oxidative and ER stress, and remodeling of extracellular matrix components.^5–7^

Significant weight loss, such as occurs after bariatric surgery, can rapidly restore insulin sensitivity, with many patients experiencing remission of T2D within days to weeks. Accordingly, there has been significant interest in assessing how weight loss affects the cellular composition and gene expression of adipose tissue. Three recent studies have described changes in human adipose tissue following bariatric surgery at single cell resolution, with all noting at least some degree of restoration of cellular composition and gene expression.^8–10^ Notably, these studies are not fully concordant on some points, with some studies noting partial retention of obesity-associated features, while others suggest a more complete restoration to the original lean state. All published studies have focused on timepoints ranging from six to >24 months after bariatric surgery, long after metabolic function has been restored.

Here we report on the results of transcriptional profiling of subcutaneous white adipose tissue (SAT) from human subjects with obesity at single cell resolution before, and at one, six, and twelve months following bariatric surgery. We track cell populations and subpopulations that change over the time course of weight loss, and identify specific alterations that may impact metabolism. Most notably, our study identifies a large number of changes that occur in the first month following bariatric surgery, including a shift among adipocytes along two separate axes, suggesting simultaneous apoptosis of old, stressed adipocytes and their replacement through new differentiation of healthy cells. Moreover, we provide evidence that the process by which new adipogenesis is enabled is via the surgery-induced loss of a hedgehog signaling axis within the adipose niche involving a specific endothelial cell subpopulation and an anti-adipogenic population of ASPCs. Taken together, our data point to an extremely rapid and dynamic shift in adipose cellular composition and function early after bariatric surgery.

## RESULTS

### A longitudinal profile of subcutaneous adipose tissue after bariatric surgery

To determine the molecular changes in adipose tissue that follow bariatric surgery, we recruited 14 people with obesity, representing both sexes and diverse ages. These subjects were naive to GLP1 receptor agonists, and underwent either Roux-en-Y gastric bypass (RYGB, n=7, 3 male, 4 female) or vertical sleeve gastrectomy (VSG, n=7, 4 male, 3 female). Abdominal SAT samples were collected during the initial surgery and at follow-up visits at one, six, and twelve months post-surgery (**Figure 1A**). All participants showed significant weight loss and metabolic improvement, including reduced Homeostasis Model Assessment of Insulin Resistance (HOMA-IR), within one month of the procedure (**Figures 1B, C** and **S1**).

**Figure 1.**
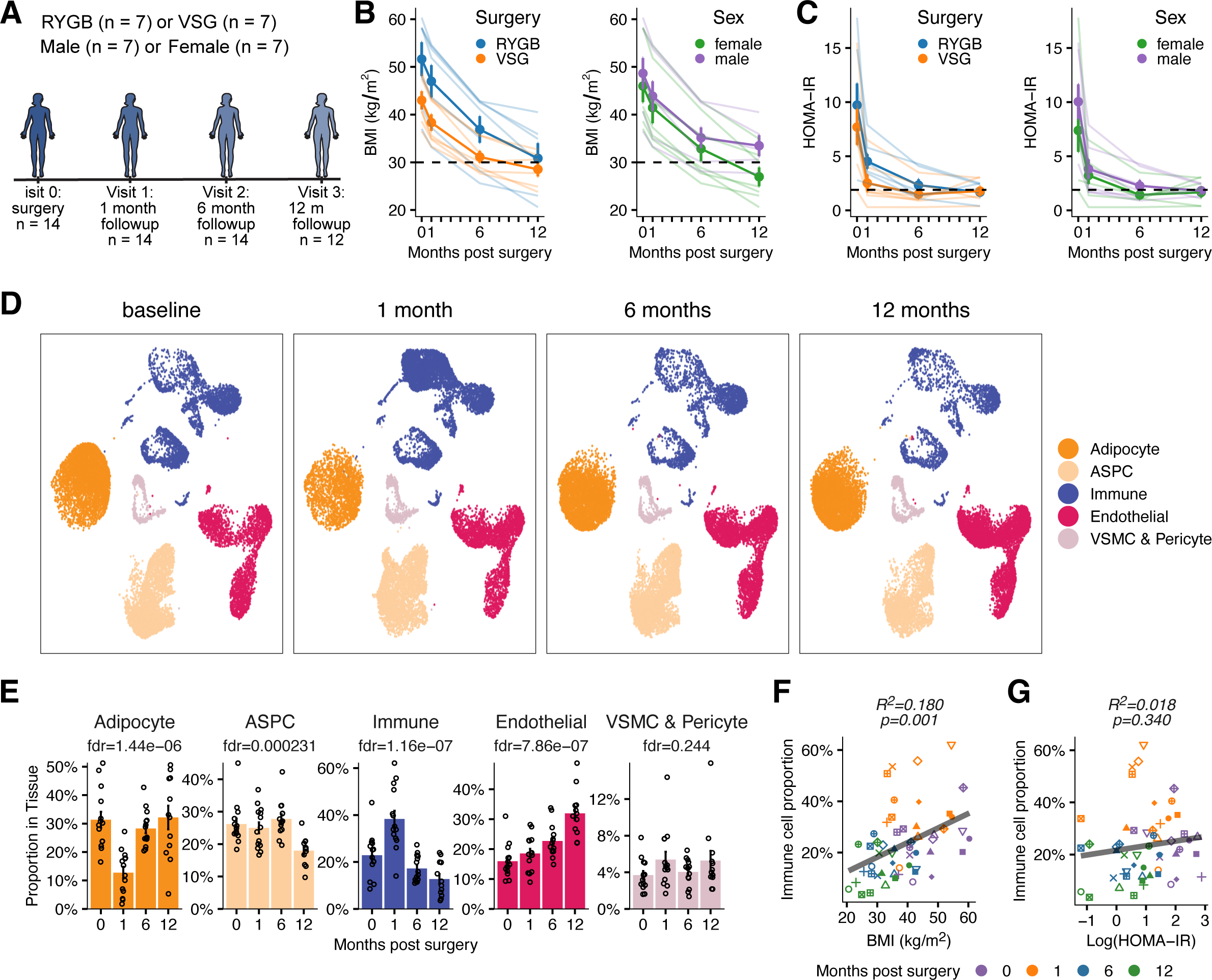
The composition of human adipose tissue changes early after bariatric surgery. (A) Overview of the study design. (B) Body mass index (BMI) of participants following surgery, grouped by procedure (left panel) and sex (right panel). Thin colored lines represent individual participants; thick lines and data points indicate aggregated results across participants. (C) Homeostatic Model Assessment of Insulin Resistance (HOMA-IR) of participants following surgery, grouped by procedure (left panel) and sex (right panel). Thin colored lines represent individual participants; thick lines and data points indicate aggregated results across participants. (D) Uniform manifold approximation and projection (UMAP) of snRNA-seq data from adipose tissues of 14 donors, split by timepoint. A total of 112,991 nuclei passed quality filtering, and plots were down-sampled to 15,997 nuclei per timepoint for visual comparison. Colors indicate major cell types. (E) Relative proportions of major cell types at designated timepoints. Data are represented as mean ± SEM. Statistical significance of variations in cell type proportions across different timepoints was assessed using the propeller method implemented in R package speckle. (F) The relationship between the proportion of immune cells and the corresponding donor’s BMI at the time of sampling. Individual participants are differentiated by point shape, and different timepoints are indicated by color. (G) The relationship between the proportion of immune cells and the corresponding donor’s HOMA-IR score at the time of sampling. Individual participants are differentiated by point shape, and different timepoints are indicated by color.

We performed snRNA-seq on the 54 available tissue samples (two subjects were lost to follow-up at the 12 month timepoint), yielding transcriptomic profiles for 112,991 nuclei after filtering for quality control; all expected cell types in human SAT were identified (**Figures 1D** and **S2A**). We found no major differences in transcriptional profiles when comparing RYGB to VSG surgery, or between men and women (**Figures S2B-E**). Accordingly, subjects were pooled in subsequent analyses in order to maximize analytical power for our primary focus on differences across post-surgical timepoints.

Adipose tissue composition underwent major changes post-surgery, with many distinctive alterations observed at the earliest time points (**Figures 1D, E)**. At the one month mark, the immune cell component nearly doubled as a proportion of all cell types relative to baseline, before declining at the 6 and 12 month time points. This was especially striking as the doubling of the immune cell population coincided with the most robust improvements in HOMA-IR. In fact, while the proportion of immune cells correlated reasonably well with BMI across all time points (R^2^ =0.180, p=0.001), there was no significant relationship between immune cells and HOMA-IR (R^2^ =0.018, p=0.340) (**Figures 1F, G**). At one month, the proportion of adipocytes declined significantly, an effect partially attributed to extensive immune cell recruitment at that time point. By six and twelve months post-surgery, adipocyte proportions were comparable to baseline, in agreement with prior studies demonstrating that adipocyte number remains constant after weight loss.^11,12^ Interestingly, the proportion of endothelial cells increased progressively following surgery. This increase was negatively correlated with both BMI and HOMA-IR, suggesting a potential link between improved adipose tissue vascularization and better overall metabolic health (**Figures S2F, G**).

As an orthogonal way to probe how these changes in adipose composition might affect systemic physiology, we connected our findings to changes in the plasma proteome using an independent dataset in which 4,151 proteins were profiled in the plasma of 35 non-diabetic adults at baseline, 1-2 months, and 6 months following RYGB.^13^ All participants achieved significant weight loss (**Figure S3A**), and their plasma profiles were consistent with improved metabolism, including elevated SHBG and IGFBP isoforms (**Figure S3B**). Moreover, levels of well-known adipokines like leptin (encoded by *LEP*) and adiponectin (encoded by *ADIPOQ*) decreased and increased, respectively, within 1-2 months (**Figure S3C**), consistent with their known behavior following weight loss. Also, in agreement with the snRNA-seq data, enrichment analysis of proteins significantly downregulated post-surgery indicated a marked decrease in inflammatory and immune responses and an increase in pathways associated with vascular development (**Figure S3D-G**).

### Recruitment of lipid-associated macrophages immediately after bariatric surgery

To further characterize post-surgical immune profile changes, we subclustered the immune cell populations and assigned cell type based on marker gene expression (**Figures 2A, B**). Most immune cell types remained stable or slowly decreased in proportion after surgery (**Figures 2C-E**). A notable exception was a macrophage subpopulation marked by the fatty acid binding protein 4 (*FABP4*) gene. These cells, previously described as lipid-associated macrophages (LAM),^14^ become significantly increased in representation one month after surgery, accounting for approximately 20% of all cells in the fat pad (**Figures 2C-E**). Histological evaluation of a representative donor revealed a striking, transient accumulation of crown-like structures surrounding adipocytes at one month post-surgery, which subsequently resolved by 6 and 12 months (**Figure 2F**). These cells, which possess a distinct capability for lipid handling, are marked by the triggering receptor expressed on myeloid cells 2 (*TREM2*) gene, as well as *IL1RN* (**Figure 2G, S4A)**. The plasma proteomic data was consistent with these results, showing that the LAM markers FABP4, TREM2 and IL1RN were transiently up-regulated in the blood at one to two months post-surgery (**Figure 2H**).

**Figure 2.**
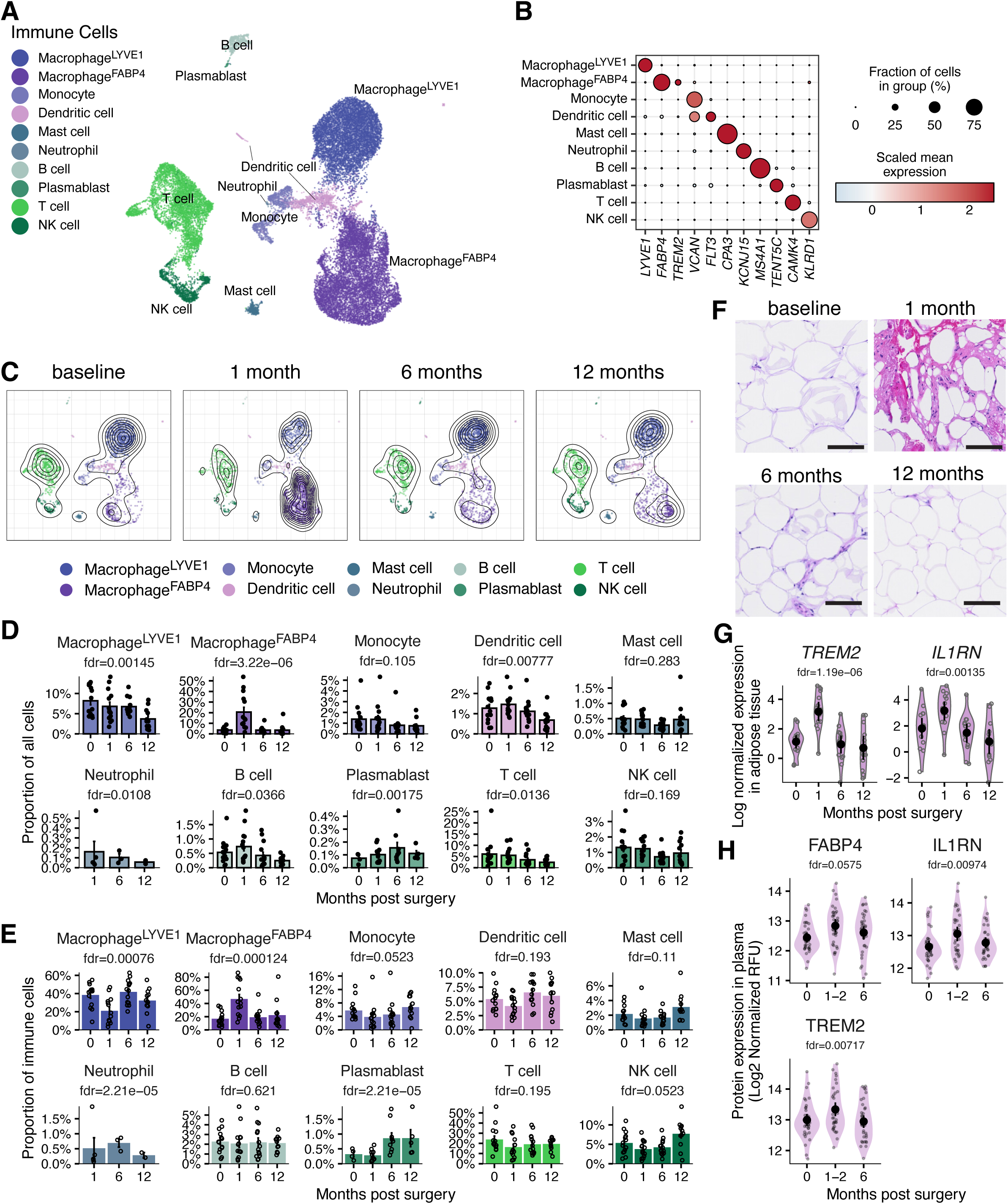
Changes in immune cell classes in SAT after bariatric surgery. (A) UMAP projection of 28,136 immune cells. (B) Expression of immune cell type markers. (C) UMAP projections of immune populations, separated by sampling timepoint. Each plot is normalized to an equal cell count (n = 1,979), with 2D density contours overlayed. (D) Relative proportions of immune cell types as a percentage of all adipose tissue cells across timepoints. Data are represented as mean ± SEM. Statistical significance of variations in cell type proportions across different timepoints was assessed using the propeller method implemented in R package speckle. (E) Relative proportions of immune cell types as a percentage of all immune cells across timepoints. Data are represented as mean ± SEM. Statistical significance of variations in cell type proportions across different timepoints was assessed using the propeller method implemented in R package speckle. (F) Representative haematoxylin and eosin staining of SATs from a single donor. Scale bar, 100 μm. (G) Expression of *IL1RN* and *TREM2* genes (associated with Macrophage^FABP4^) in SAT following surgery. Data are presented as log2-transformed counts per million. Statistical significance was determined via the Quasi-Likelihood F-test. (H) Levels of IL1RN and TREM2 proteins in plasma after bariatric surgery (n = 35 subjects). Repeated measures ANOVA was used to evaluate statistical significance. Data distribution shown via violin plots (mean ± 95% bootstrapped CI).

### Increased vascularization accompanies the improvement of the metabolic profile

Our analysis identified all of the major vascular cell types in adipose tissue, including arterial, capillary, and venous blood endothelial cells, lymphatic endothelial cells, smooth muscle cells, and pericytes (**Figures 3A, B**). We noted progressive increases in the proportion of endothelial cells over the year following surgery, while lymphatic endothelial cells remained constant (**Figures 1E, 3C**), a finding corroborated by immunostaining (**Figure 3D**). As mentioned, this aligns with the plasma proteomic data, which showed an up-regulation of endothelial development pathways among circulating plasma proteins following bariatric surgery (**Figures S3F, G**). Notably, one subpopulation, denoted as Arterial^PTPRJ^ endothelial cells, decreased significantly after surgery (**Figures 3E, F**; discussed further below). The relative proportions of other arterial, venous, and capillary cells remain largely unchanged within the context of the overall expanding vasculature.

**Figure 3.**
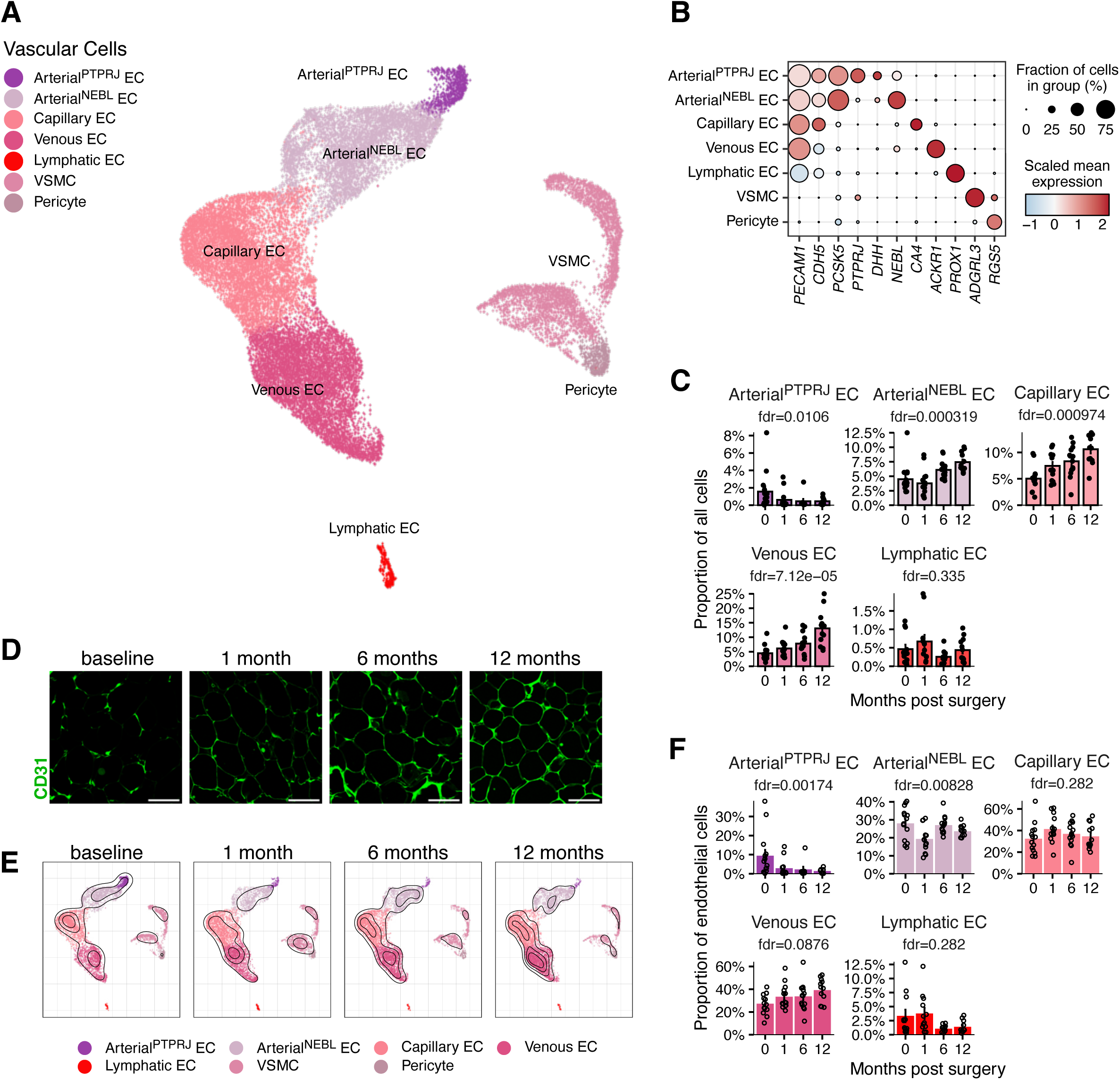
Changes in vascular cell classes in SAT after bariatric surgery. (A) UMAP projection of 27,874 vascular cells. (B) Expression of vascular cell type markers. (C) Relative proportions of vascular cell types as a percentage of all adipose tissue cells across timepoints. Data are represented as mean ± SEM. Statistical significance of variations in cell type proportions across different timepoints was assessed using the propeller method implemented in R package speckle. (D) Representative immunofluorescence staining for CD31 expression. Scale bars, 100 μm. (E) UMAP projections of vascular cell populations, separated by sampling timepoint. Each plot is normalized to an equal cell count (n = 5,844), with 2D density contours overlayed. (F) Relative proportions of vascular cell types as a percentage of all vascular cells across timepoints. Data are represented as mean ± SEM. Statistical significance of variations in cell type proportions across different timepoints was assessed using the propeller method implemented in R package speckle.

### Adipocyte subpopulations shift dramatically soon after bariatric surgery

The spatial distribution of all adipose tissue cell types in the UMAP embedding suggested significant changes in the adipocyte profile following bariatric surgery (**Figure 1D**). To more finely characterize these changes, adipocytes were clustered into 10 subpopulations with distinct transcriptional profiles (**Figures 4A-C, S4B, C**). At baseline, Ad^AKR1C2^ and Ad^CARD18^ comprised the majority of adipocytes, but their representation significantly decreased by one month post-surgery and remained low thereafter (**Figure 4D**). Concurrently, the Ad^VSTM4^ population progressively expanded, comprising 59.2% of adipocytes at twelve months post-surgery (**Figure 4D**). Pathway analysis of markers indicates that Ad^CARD18^ likely represents ‘stressed’ adipocytes, with enrichment of pathways like the unfolded protein response (**Figure 4E**). Conversely, Ad^VSTM4^ markers are enriched in adipogenesis and glycerolipid biosynthesis pathway genes, suggesting that these represent healthier adipocytes (**Figure 4E**). Collectively, these data support the idea that adipocytes move from a biochemically stressed state in obesity to a more metabolically healthy profile by 6-12 months after surgery, consistent with prior observations.^8-10^

**Figure 4.**
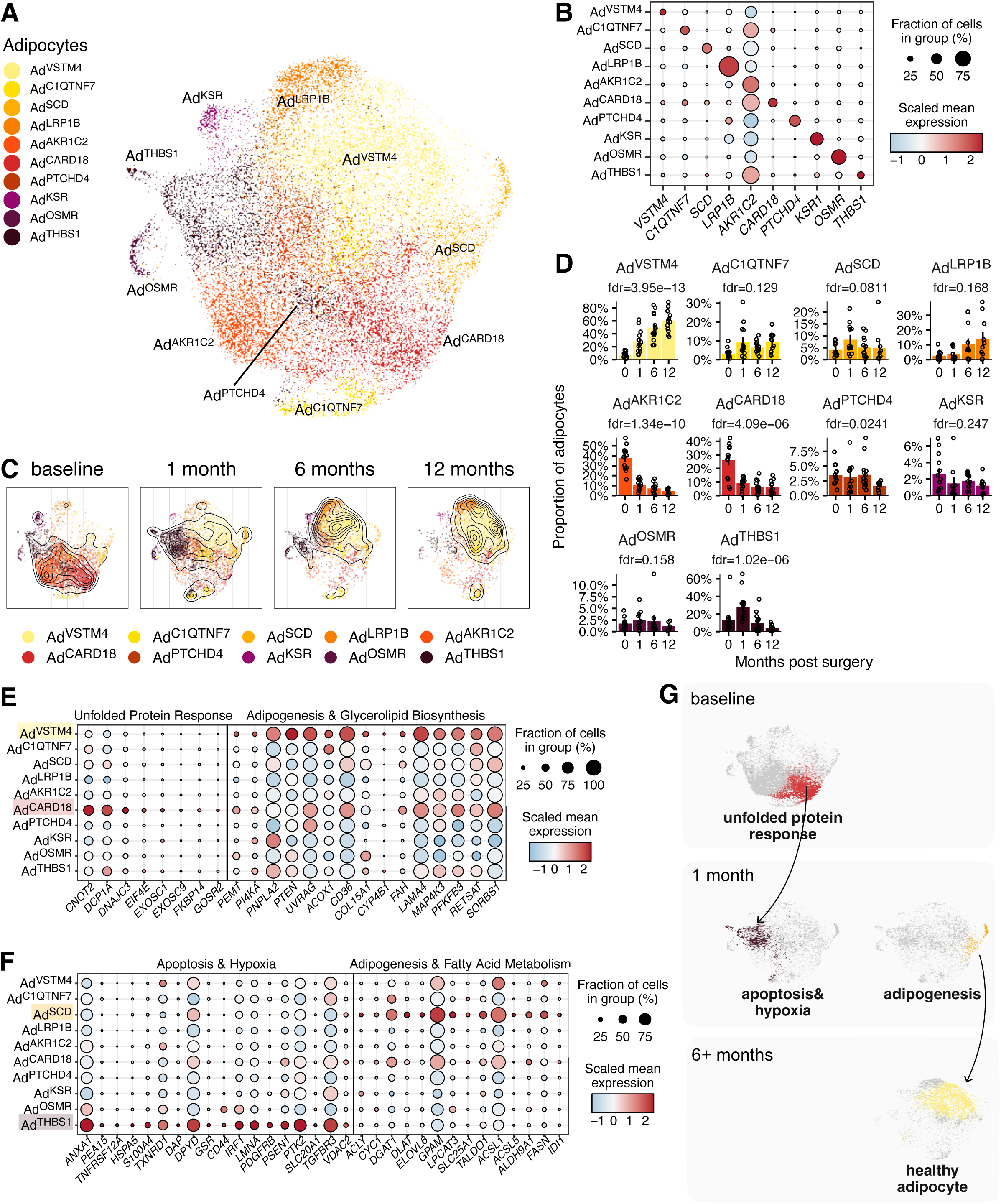
Changes in adipocyte subtypes in SAT after bariatric surgery. (A) UMAP projection of 29,033 adipocytes. (B) Expression of adipocyte subtype markers. (C) UMAP projections of adipocyte populations, separated by sampling timepoint. Each plot is normalized to an equal cell count (n = 4,183), with 2D density contours overlayed. (D) Relative proportions of adipocyte subtypes as a percentage of the overall adipocyte population across timepoints. Data are represented as mean ± SEM. Statistical significance of variations in cell type proportions across different timepoints was assessed using the propeller method implemented in R package speckle. (E) Expression of genes associated with the “unfolded protein response”, “adipogenesis”, and “glycerolipid biosynthesis” pathways in each adipocyte subtype. (F) Expression of genes associated with the “apoptosis”, “hypoxia”, “adipogenesis” and “fatty acid metabolism” pathways in each adipocyte subtype. (G) Proposed model of adipocyte turnover dynamics in SAT following bariatric surgery.

We noted the transient appearance of two subpopulations, Ad^THBS1^ and Ad^SCD^, both of which showed highest representation at the one month timepoint (**Figure 4D**). Ad^THBS1^ adipocytes, which make up 27.4% of adipocytes at one month after surgery, exhibit markers enriched for ‘apoptosis’ and ‘hypoxia’ genes, suggesting that these represent adipocytes that may be in the process of being cleared from the tissue (**Figure 4F**). This is consistent with our observation that lipid-associated macrophages (Macrophage^FABP4^) rise at one month post-surgery, as these cells have been implicated in clearing dead and dying adipocytes (**Figure 2C, D** and ^15^). In contrast, Ad^SCD^ cells are enriched for markers of adipogenesis and fatty acid metabolism (**Figure 4F**). Given that the overall adipocyte proportion remains stable post-surgery (**Figure 1E**)^11,12^, we speculate that Ad^SCD^ may represent newly developed adipocytes, although there is, to our knowledge, no specific transcriptional marker for a “young” adipocyte. Collectively, the contrasting profiles of Ad^THBS1^ (apoptosis/hypoxia) and Ad^SCD^ (adipogenesis/metabolism), alongside the elimination of Ad^CARD18^ (stressed) and establishment of Ad^VSTM4^ (healthy), provide evidence supporting a working model of adipocyte turnover following bariatric surgery in which at least some older, stressed adipocytes undergo cell death and are replaced by new, healthy adipocytes (**Figure 4G**).

### Adipogenesis is detected after bariatric surgery

To further assess whether adipogenesis is heightened in the early phases of weight loss, we looked at the adipose stromal and progenitor cell (ASPC) population, which comprises a heterogeneous mesenchymal population with varying degree of adipogenic potential. Through subclustering, marker gene analysis and reference mapping, we identified known ASPC subtypes, including *DPP4*^+^ early progenitor clusters (ASPC^DPP4+,ABCB5+^ and ASPC^DPP4+,ABCB5-^), *PPARG*^+^ committed preadipocyte clusters (ASPC^PPARG+,POSTN-^ and ASPC^PPARG+,POSTN+^), as well as ASPC^PTCH2^, which most closely resembles a population of anti-adipogenic regulatory cells called Aregs (**Figures 5A-D, S4D, E**).^16,17^

**Figure 5.**
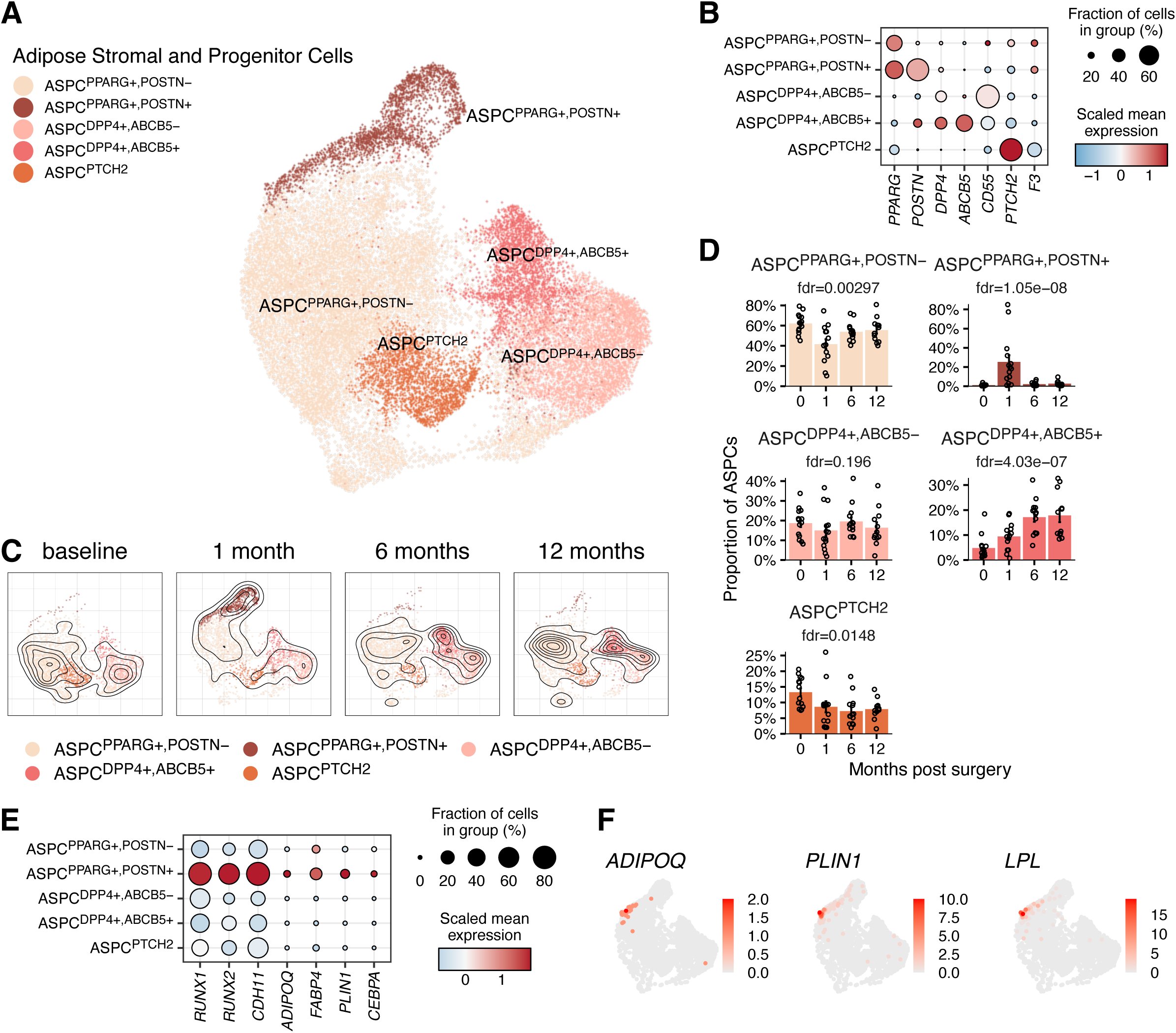
Subclustering of adipose stromal and progenitor cells (ASPCs) (A) UMAP projection of 27,948 ASPCs. (B) Expression of ASPC cell type markers. (C) UMAP projections of ASPC populations, separated by sampling timepoint. Each plot is normalized to an equal cell count (n = 2,890), with 2D density contours overlayed. (D) Relative proportions of ASPC subtypes as a percentage of the overall ASPC cell population across timepoints. Data are represented as mean ± SEM. Statistical significance of variations in cell type proportions across different timepoints was assessed using the propeller method implemented in R package speckle. (E) Expression of selected ASPC^PPARG+,POSTN+^ markers. (F) UMAP projection of ASPCs colored by expression of adipocyte markers.

Notably, ASPC^PPARG+,POSTN+^ is a transient subpopulation observed almost exclusively at one month post-surgery (**Figure 5C, D**). These cells exhibit a transcriptional profile that is distinctive from other ASPC populations reported in this study and in the literature.

ASPC^PPARG+,POSTN+^ cells are characterized by the expression of transcription factors, including *RUNX1* and *RUNX2*, that are known for their role in mesenchymal development, as well as the adipogenic transcription factors *CEBPA* and *PPARG* (**Figure 5E**).^18–23^ Furthermore, a subset of ASPC^PPARG+,POSTN+^ cells distinctively express adipocyte marker genes such as *ADIPOQ*, *PLIN1*, and *LPL*, indicating they are differentiating progenitors actively acquiring adipocyte identity (**Figures 5E, F**). Analysis of the lineage trajectory shows that our data recapitulates the known ontogeny of adipogenesis, with *DPP4*^+^ cells representing the earliest ASPCs, occurring earlier than ASPC^PPARG+,POSTN-^ preadipocytes, which then diverge into non-adipogenic ASPC^PTCH2^ cells (Aregs) and ASPC^PPARG+,POSTN+^ preadipocytes, the latter of which directly precedes the differentiated adipocyte state (**Figure S4F**). Collectively, these analyses corroborate the suggestion that active progenitor differentiation is most prominent at one month post-surgery.

### Obesity upregulates anti-adipogenic hedgehog signaling, resolved with weight-loss

We next sought to identify processes that might account for increased adipogenesis following weight loss surgery. We noted that ASPC^PTCH2^ cells were significantly reduced in representation post-surgery (**Figure 5D**). This population most closely resembles cells identified by the Deplancke and Wolfrum group as adipogenic regulatory cells, or Aregs, which act to reduce adipogenesis (**Figure S4E**).^17^ Review of ASPC^PTCH2^ markers and enrichment analysis revealed that they express genes associated with the hedgehog signaling pathway (**Figures 6A, B**), which is known for its role in developmental patterning and which has been shown to exert a strong anti-adipogenic effect.^24–27^ It is known that adipogenesis is diminished in the obese state^28^; the relatively high proportion of ASPC^PTCH2^ cells could explain this, and their rapid decline following bariatric surgery might enable new adipogenesis.

**Figure 6.**
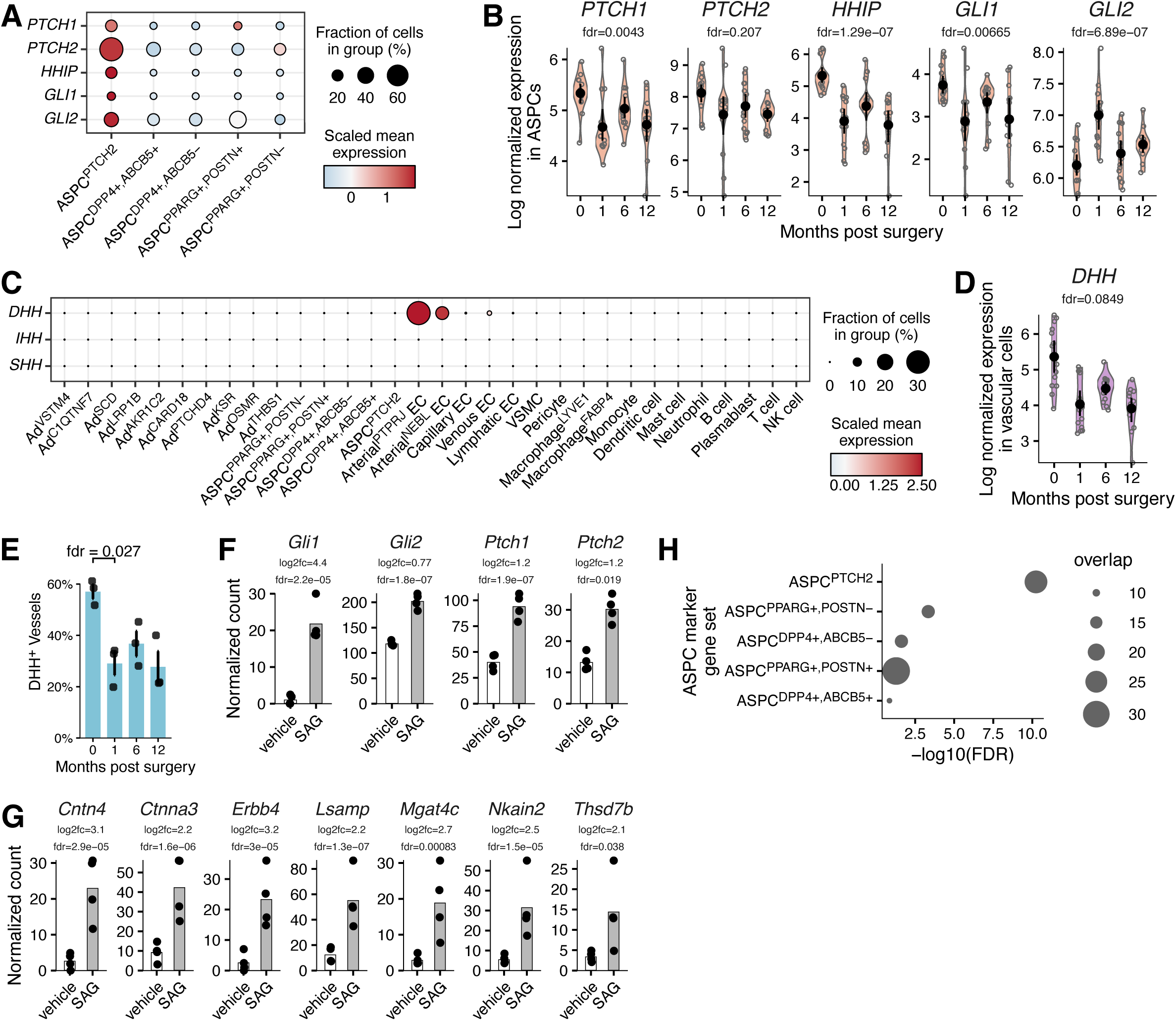
Anti-adipogenic hedgehog signals diminish after bariatric surgery. (A) Expression of hedgehog pathway genes in ASPC subtypes. (B) Expression of hedgehog pathway genes in SAT over the course of this study. Data are presented as log2-transformed counts per million. Statistical significance was determined via the Quasi-Likelihood F-test. (C) Expression of hedgehog ligands in all SAT cell subtypes. (D) Expression of *DHH* across timepoints. Data are presented as log2-transformed counts per million. Statistical significance was determined via the Quasi-Likelihood F-test. (E) Quantification of the percentage of DHH^+^ vessels performed on adipose tissues from three individuals at four post-surgical time points. Each data point represents the aggregate average calculated from five images acquired from the corresponding tissue sample. Bar plot data are represented as mean ± SEM. Repeated-measures ANOVA was used to assess statistical significance. (F) Expression of hedgehog pathway genes in primary mouse stromal vascular fraction (mSVF) treated with 200 nM of SAG or vehicle for 24 hours (n=3 per treatment condition). Bar plot data are represented as mean. Statistical significance was assessed using the Wald test implemented in the DESeq2 R package. (G) Expression of ASPC^PTCH2^ markers in mouse SVF treated with 200 nM of SAG or vehicle for 24 hours (n=3 per treatment condition). Bar plot data are represented as mean. Statistical significance was assessed using the Wald test implemented in the DESeq2 R package. (H) Enrichment analysis using a hypergeometric test performed on genes differentially upregulated in mSVF treated with 200 nM SAG, compared to vehicle treatment. The gene sets used for this analysis were based on markers for ASPC subtypes.

ASPC^PTCH2^ cells have the machinery to respond to hedgehog proteins, but they do not produce the ligand themselves. We therefore looked across other cell types in the fat pad to see where hedgehog family members might be generated. Among the hedgehog ligands, only desert hedgehog (*DHH*) is expressed in adipose tissue, relatively specifically by a subtype of arterial endothelial cells, Arterial^PTPRJ^ EC (**Figure 6C**). Following bariatric surgery, the proportion of this endothelial population and the expression of *DHH* both decreased, temporally correlating with the reduction in ASPC^PTCH2^ (**Figures 5D**, **6D, E** and **S5A**). Of note, Areg cells have been characterized as a perivascular stromal cell type,^17^ consistent with current belief that hedgehog signaling acts in a paracrine manner over a short range. These findings are consistent with a model where direct signaling from endothelial *DHH*, specifically from the Arterial^PTPRJ^ EC population, promotes ASPC^PTCH2^/Areg identity and/or function.

While our data highlights unique features of human adipose tissue remodeling, obtaining sufficient primary human ASPCs from this clinical cohort for mechanistic studies was not feasible. To test the idea that hedgehog signal promotes ASPC^PTCH2^/Areg identity, we profiled the transcriptomes of primary mouse stromal vascular fraction (SVF) treated with either Smoothened receptor agonist (SAG) or recombinant human desert hedgehog (DHH), followed by bulk RNA sequencing. Treatment with these agonists up-regulated the expression of both hedgehog target genes and ASPC^PTCH2^ markers (**Figures 6F, G** and **S5B, C**). This effect on ASPC^PTCH2^ markers shows some specificity, in that genes that mark other ASPC populations are not up-regulated by hedgehog agonists (**Figures 6H** and **S5D**). Taken together, these data are consistent with a model in which endothelial-Areg hedgehog paracrine axis suppresses adipogenesis in the obese state, and becomes rapidly attenuated after bariatric surgery.

### Senescence profiles improve following bariatric surgery, but follow different time courses in different cell types

Senescence of many cell types in adipose tissue, including adipocytes, has been posited to drive metabolic deterioration in obesity, and recent reports suggest that bariatric surgery reduces adipose senescence.^9,29,30^ We sought to determine whether this held true in our dataset. We first noted, however, that “senescence” gene sets differ greatly between published studies. We therefore took four recent gene sets purported to relate to senescence,^9,31–33^ and focused on the nine genes that appeared in at least three of them (*CDKN1A*, *CDKN2A*, *TP53*, *CDK2*, *CDK6*, *ETS2*, *IGFBP7*, *IL1A*, and *IL6*) (**Figure S6A**). Adipocytes showed a steady and sequential decline in the core senescence marker gene expression across all timepoints. In contrast, in ASPCs, vascular, and immune cells, many senescence genes are actually elevated at the one month timepoint, before falling again at the 6 and 12 month timepoints (**Figures S6B, C**).

### Lack of concordance between mouse and human adipose profiles after bariatric surgery

We have previously published a single cell analysis of white adipose tissue after VSG in mice,^34^ which included early timepoints (10 and 35 days post-surgery), enabling us to compare the effect of bariatric surgery across species, with two important caveats. First, the mouse data do not represent serial samples on the same individual, as was done in the human. Second, by 35 days, mice with VSG had regained much of their lost weight, although they still weighed significantly less than the 35 day sham-operated control mice (**Figure 7A**). Spontaneous weight regain after VSG in mice has been noted by others as well.^35^ With these caveats in mind, we identified similarities and differences in adipose tissue changes.

**Figure 7.**
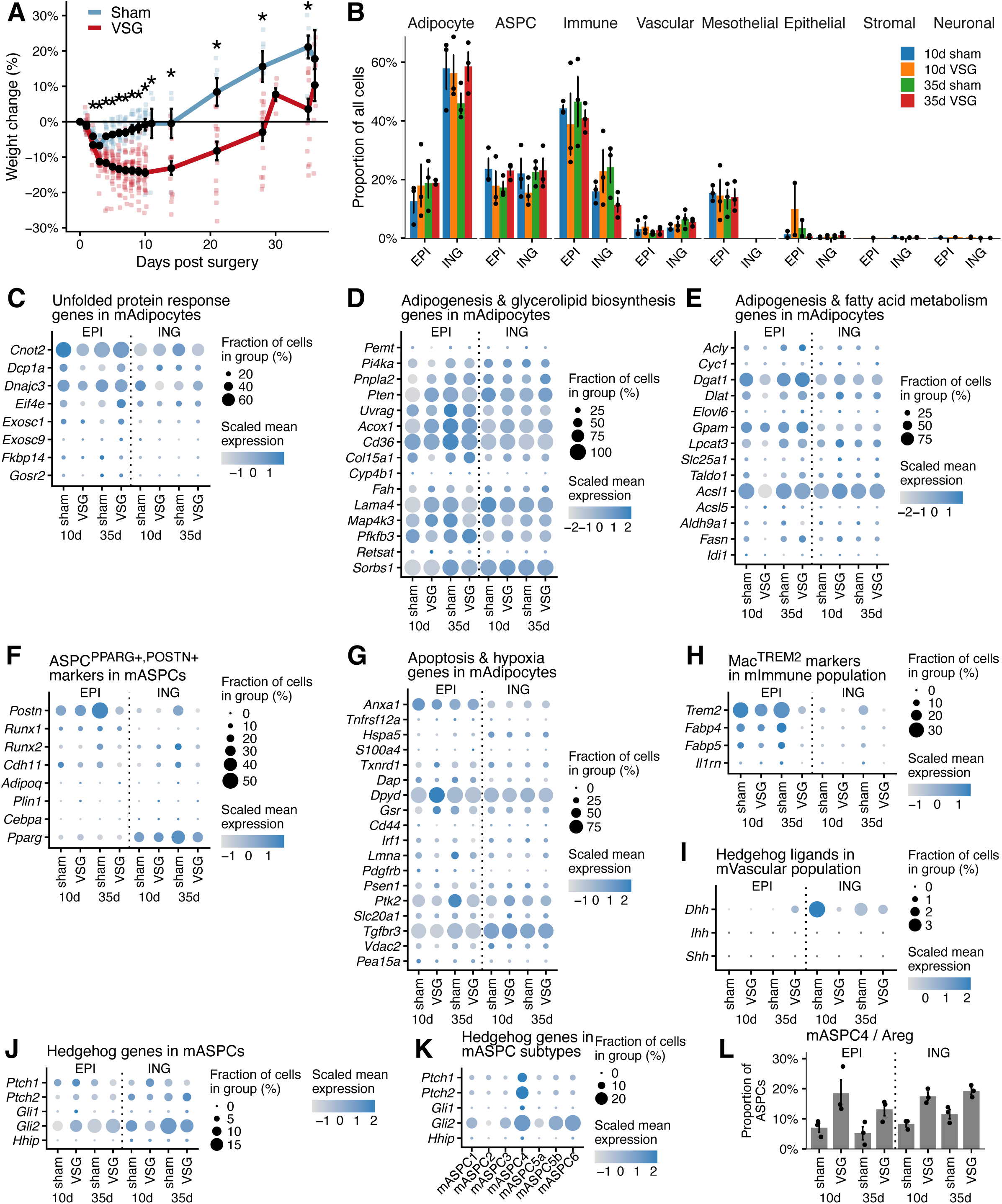
Findings in human adipose tissue post-bariatric surgery are only partially mirrored in the bariatric surgery mouse model. (A) The percent weight changes in mice following either VSG or sham surgeries, with data sourced from Emont, et al. Error bars represent the standard error. Statistical significance was assessed using a student’s *t*-test to compare the body weights of the VSG and sham groups, with correction for multiple comparisons applied via a false discovery rate (FDR). *FDR < 0.05. (B) Relative proportions of major cell types in mouse epididymal (EPI) or inguinal (ING) adipose tissue at designated timepoints after VSG. (C) Expression of the unfolded protein response related genes that were observed in human Ad^CARD18^, in mouse adipocytes after VSG. (D) Expression of adipogenesis and glycerolipid biogenesis related genes that were observed in Ad^VSTM4^, in mouse adipocytes after VSG. (E) Expression of adipogenesis and fatty acid metabolism related genes that were observed in Ad^SCD^, in mouse adipocytes after VSG. (F) Expression of ASPC^PPARG+,POSTN+^ markers in mouse ASPCs after VSG. (G) Expression of apoptosis and hypoxia genes, observed in Ad^THBS1^, in mouse adipocytes after VSG. (H) Expression of Mac^TREM2^ markers in the immune cells of mouse adipose tissue after VSG. (I) Expression of Hedgehog ligands in the vascular cells of mouse adipose tissue after VSG. (J) Expression of Hedgehog pathway genes in mouse ASPCs after VSG. (K) Expression of Hedgehog pathway genes in mouse ASPC subtypes. (L) Relative proportion of mASPC4/Areg in EPI or ING adipose tissues at designated timepoints after VSG.

As expected, some of the changes that develop over 6-12 months in humans are not readily apparent in this short-term mouse experiment. For example, the overall proportion of vascular and immune cells remained unchanged after VSG in mice, in both inguinal and epididymal depots (rough orthologs of human SAT and visceral adipose tissue, respectively)(**Figure 7B**). Similarly, while genes associated with the unfolded protein response are prominent in adipocytes from obese mice, there is a mixed response to VSG, with some diminishing (e.g., *Cnot2*, *Dnajc3*) while others increase (e.g., *Exosc1*), often in a depot-specific manner (**Figure 7C**). The Ad^VSTM4^ markers, indicative of healthy, “destressed” adipocytes, are also not observed after VSG (**Figure 7D**).

The mouse data at the studied time points and depots do not, however, suggest that bariatric surgery induces an adipocyte turnover process analogous to what we observed in humans. Specifically, we observed no up-regulation of adipogenesis genes (indicated by Ad^SCD^ or Ad^VSTM4^ markers) or the emergence of an ortholog of ASPC^PPARG+,POSTN+^ cells (**Figure 7D-F**). Similarly, while a few selected apoptosis and hypoxia genes that define human Ad^THBS1^ are expressed in adipocytes of obese mice, we do not see a response suggesting that apoptosis increases in either depot at the 10 day or 35 day timepoint (**Figure 7G**). This is concordant with the absence of Macrophage^FABP4^/LAM expansion in the VSG group (**Figure S7H**), which may be required to police the adipose niche following adipocyte apoptosis.

Similar to the human data, we note that *Dhh* is the dominant hedgehog isoform in mouse adipose tissue, and that it is expressed in endothelial cells. *Dhh* expression is high in the inguinal adipose tissue of obese mice, and it diminishes after VSG (**Figure 7I**). Expression of hedgehog signaling genes in the ASPC population, however, shows a more mixed pattern with reduction of some genes (e.g., *Gli2*) and moderate increases in *Ptch2* (**Figure 7J**). Contrary to what we noted in humans, the Areg population in mouse adipose tissue (identified as mASPC4 in our prior mouse work) increases with weight loss in all depots and time points tested (**Figure 7K, L**).

## DISCUSSION

In this study, we provide a longitudinal characterization of the cellular changes in human SAT during bariatric surgery-induced weight loss. By tracking the same patient cohort over a one-year period, we were able to observe the tissue as it transitioned from a state of obesity-associated metabolic stress to a post-weight-loss healthier steady state. Our data demonstrate significant shifts in the transcriptional identity and proportions of nearly every major cell type within the adipose niche. While other single-cell studies have described the adipose landscape in response to surgically-induced weight loss,^8–10^ our study is unique in capturing the early one month post-operative time point. This window enabled us to see early changes during a phase of remodeling that correlates temporally more closely with improvements in metabolic function. One obvious illustration of the power of this approach is our observation that the long-term decrease in immune cell infiltration reported by others (and seen by us at later timepoints) is preceded by a transient upregulation of Macrophage^FABP4^/LAMs at one month. As discussed below, we speculate that this early response may be essential for the clearance of apoptotic, lipid-laden adipocytes and the initiation of healthy tissue turnover. Lipid-associated macrophages have been linked to obesity-associated insulin resistance^36^; in contrast, we note that the transient elevation in these cells in SAT following bariatric surgery was temporally associated with profoundly improved HOMA-IR.

There is also a complex relationship between body weight and adipose vascularity. In general, obesity is considered to drive relative hypoxia in the fat pad, which raises VEGF expression in adipocytes and in plasma.^37–39^ Whether this leads to increased vascularity in obesity is unclear, however. Single cell data from lean and obese human adipose tissue suggest a lower proportion of endothelial and other vascular cell types in samples from obese vs. lean subjects; this is in line with deconvoluted bulk RNA-seq data from the METSIM cohort.^40,41^ In contrast, little is known about the response of the adipose vasculature to weight loss. Lower vascular density has been correlated with more robust weight loss after bariatric surgery, but this study used samples collected at the time of surgery and did not follow vascularity over time.^42^ A recent study by the Mandrup group indicates that vascularity increases in SAT after bariatric surgery,^10^ which we also observe.

In both humans and mice, adipocytes in the obese state exhibit signs of stress, with up-regulation of genes associated with hypoxia,^43^ inflammation,^44^ and insulin resistance,^45^ seen also in our baseline samples. By the 6-12 month post-surgical timepoints, the gene expression pattern of adipocytes has shifted dramatically, with up-regulation of fatty acid metabolic genes and other hallmarks of healthy adipocytes. What accounts for the presence of these healthy adipocytes after weight loss? Do they represent older adipocytes that have ‘de-stressed’, or are they new adipocytes that differentiated following the surgical intervention? If the latter, what has become of the older, stressed fat cells? Our transcriptional study cannot definitively answer these questions, but at one month following surgery, we note that the adipocyte compartment exhibits evidence of two separate processes: apoptosis of older, stressed adipocytes and the simultaneous formation of new, healthy fat cells.

Healthy adipocytes are relatively resistant to apoptosis, in part because of insulin action, and also because they express many anti-apoptotic factors.^46^ Obesity, however, is associated with enhanced apoptosis of adipocytes, with dead and dying fat cells surrounded by a corona of pro-inflammatory macrophages called a crown-like structure.^15,47^ Interestingly, blockade of adipocyte apoptosis through targeted deletion of caspases or other proapoptotic factors prevents excess weight gain on high fat diet and reduces adipose inflammation and insulin resistance.^47–50^ Our data suggest that rather than relieve this apoptotic state, the early stages of weight loss are characterized by enhanced adipocyte death. One subpopulation of adipocytes (Ad^THBS1^) is notable for its expression of pro-apoptotic genes, including *VDAC2*,^51,52^ *TGFBR3*,^53^ *ANXA1*,^54^ *DAP*,^55^ and *IRF1*.^56^ These cells represent approximately 15% of subcutaneous adipocytes at baseline, and then increase in abundance after surgery, accounting for roughly 30% of adipocytes at one month. They then diminish over time, representing <5% of adipocytes by 1 year post-surgery. These data suggest that one possible fate of the stressed adipocyte, even after the energy imbalance that caused obesity is rectified, is death. This is consistent with the increased number of Macrophage^FABP4^/LAMs at one month, which may be required to clean up cellular debris and free lipid following apoptosis.

In both rodents and humans, the number of small adipocytes increases after weight loss.^10,57–59^ We speculate that some of this reflects delipidation of larger fat cells, but there may also be a role for new adipogenesis. Unfortunately, there is no definitive gene expression marker for a “young” adipocyte. Nevertheless, at 1 month post-surgery we note the appearance of a subpopulation of adipocytes marked by *SCD*. These cells express genes characteristic of adipogenesis. By the 6 month timepoint, these cells have largely returned to their lower baseline levels. Concordant with the appearance of these adipocytes, there is the emergence of a population of ASPCs (*PPARG+*,*POSTN+*) that express adipocyte genes such as *PLIN1*, *ADIPOQ*, and *LPL*. Taken together, our data support the notion that there is a wave of new adipogenesis at the one month timepoint that dissipates before 6 months.

Our data are consistent with a small body of literature suggesting that precursor cells isolated from individuals with obesity exhibit reduced adipogenic potential.^60,61^ This diminished adipogenic capacity has been proposed to contribute to the hypertrophy of existing adipocytes; hypertrophic adipocytes are in turn believed to be most active in promoting the metabolic dysfunction associated with obesity.^62^ Moreover, adipose precursor cells isolated after weight loss (induced by dietary restriction) demonstrate improved adipogenic potential *in vitro*.^63^

Mechanistically, how might adipogenesis be repressed in obesity, and then reactivated after weight loss? Our attention was drawn to a population of ASPCs known as Aregs (identified as ASPC^PTCH2+^ in our study), which exert anti-adipogenic actions on neighboring cells,^17^ and which decrease in abundance shortly after surgery. Notably, these cells are characterized by expression of several hedgehog receptors and signaling intermediates. Moreover, we show that *in vitro* treatment of mouse ASPCs with hedgehog agonists promotes Areg identity. We acknowledge the inherent limitation of using a murine *in vitro* system to validate a pathway identified in human tissue, particularly given that our *in vivo* mouse VSG model does not perfectly mirror the Areg population dynamics observed in humans. Nevertheless, the conserved capacity of these progenitor cells to upregulate Areg markers in response to hedgehog ligands *in vitro* provides plausible mechanistic support for the endothelial-Areg signaling axis proposed by our human transcriptomic data. Hedgehog signaling is recognized to inhibit white adipocyte differentiation in flies,^26,64^ mice,^26^ and human cells^65^; this effect is believed to involve direct actions on precursor cells, at least in culture. While we do see some hedgehog receptor expression on adipose progenitor (in particular ASPC^PPARG+POSTN-^) cells, this is dwarfed by expression on ASPC^PTCH2+^ Aregs. While Aregs respond to hedgehog signaling, they do not produce any of the recognized hedgehog ligands. Indeed, the DePlancke and Wolfrum groups identified several non-hedgehog paracrine factors (e.g., *Rtp3*, *Spink2*, *Fgf12* and *Vit*) as possible mediators of the anti-adipogenic action of Aregs.^17^

Where then, is hedgehog produced in the human adipose niche? Our data show that only one hedgehog ligand, desert hedgehog (DHH), is found in human adipose tissue, and that it is expressed exclusively by blood endothelial cells, specifically, a unique subpopulation of endothelial cells identified as Arterial^PTPRJ^ ECs. These cells rapidly decline following bariatric surgery, accompanied by reductions in *DHH* mRNA levels and immunostaining. In other contexts, DHH is involved in the response to injury.^27,66^ Specifically, muscle injury triggers the activation of fibro-adipogenic progenitors, wherein DHH is upregulated to repress fat deposition and improve muscle healing.^27^ In this light, DHH expression in obesity may represent a maladaptive injury response that prevents healthy adipose tissue homeostasis. Alternatively, DHH expression and signaling may be responding to local hypoxia in obese SAT, as hedgehog has been implicated in angiogenesis and the development of arterial endothelial cells.^67–69^ Of note, Areg cells localize to the perivascular space,^17^ consistent with the known range of hedgehog signaling as a short-range paracrine mechanism.^25,70,71^

Prior reports suggest that senescence is a driver of metabolic dysfunction in obesity, and a recent study indicates that the expression of senescence-associated genes becomes attenuated after weight loss surgery.^9^ Our data supports this idea, but a few points bear mentioning. First, there is little consensus among published gene sets about which markers truly represent markers of senescence. We tried to mitigate this issue by focusing only on genes that were included in at least three out of four published gene sets. Second, all major cell types in SAT express these ‘consensus’ senescence markers at high levels in the obese state, with diminution occurring over the 12 month course of the study period. However, only adipocytes show a stepwise reduction in the expression of these genes - the other major cell types (ASPCs, vascular, and immune cells) all display elevated senescence gene expression at the one month timepoint. It is unclear why this should be so, but one possible explanation is that these cells are exposed to the contents of apoptotic adipocytes, which induce metabolic stress and drive senescence.

We recently performed an analysis of adipose tissue composition following VSG in mice,^34^ allowing us to draw cross-species comparisons, with caveats discussed earlier. Of note, we did not detect changes in the relative amounts of vascular or immune cells in the WAT of mice after either VSG or semaglutide-mediated weight loss. This may reflect true differences between species, or perhaps the much shorter time course assessed in the mouse vs. human studies (1 month vs. 6-12 months). When assessing alterations noted at 1 month in humans, we see that there are some common features, notably elevated expression of *Dhh* in endothelial cells at baseline, which goes down following surgery in both species. We do not, however, see evidence of apoptosis of existing adipocytes, nor differentiation of new adipocytes in the first month after surgery, nor reductions in Areg levels. One interesting possibility is that failing to purge the adipose niche of large, stressed adipocytes might predispose to weight regain. Further experimentation is required to delineate why these differences exist and whether they have implications for surgical outcomes in either species.

### Limitations of the Study

Our study has several limitations. First, even though we report data from the earliest post-surgical time point yet, metabolic function had already improved significantly. Our study thus lacks the resolution to determine the precise temporal relationship between the changes in cellular composition and metabolic benefit. Second, our study was limited to SAT, due to its accessibility in an outpatient setting. We do not know whether the changes we observe in SAT are reflected in other depots, particularly omental adipose tissue. Third, a comparator group that lost weight without surgery, such as through lifestyle intervention or GLP1R agonists, would have been interesting. Lifestyle changes tend to reduce weight only a small amount and less durably, however. Finally, perhaps the most important limitation is that we can only draw inferences about the biological processes actually occurring in the adipose tissue after weight loss from changes in gene expression. There is no way, for example, to measure adipogenesis in real time in living human subjects.

## ACKNOWLEDGEMENTS

This work was supported by NIH RC2 DK116691 to EDR and NIH K01 DK134806 to MPE. We thank the Functional Genomics and Bioinformatics Core at BIDMC (supported by P30 DK135043). We thank members of the Rosen lab for helpful discussions.

## AUTHOR CONTRIBUTIONS

Experimental Design: MPE, EDR

Experimental Execution: ZYL, MPE, GPW, AG, AE, ZZ, WL, LTT

Sample Collection: WG, AC, NBK, RJS

Data Analysis: ZYL, MPE, GPW, CL, SN, JJL, LLJ, REG, EDR

Manuscript Preparation: ZYL, MPE, GPW, EDR

Supervision and Funding: EDR

## SUPPLEMENTAL FIGURES

**Figure S1.**
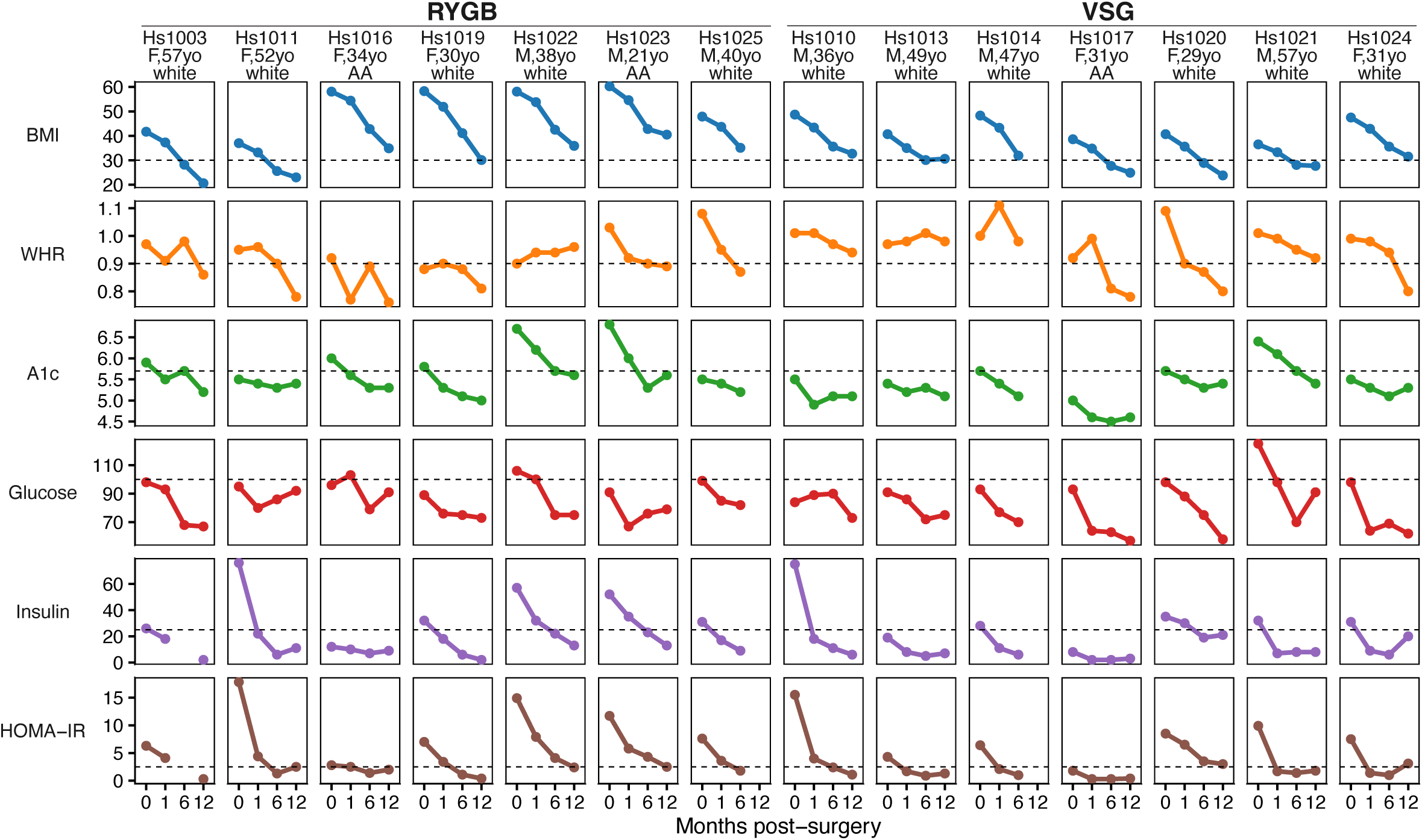
Demographic characteristics and metabolic characterization of bariatric surgery weight loss study participants across timepoints, related to main Figure 1. F = female, M = male, yo = year-old, AA = African American.

**Figure S2.**
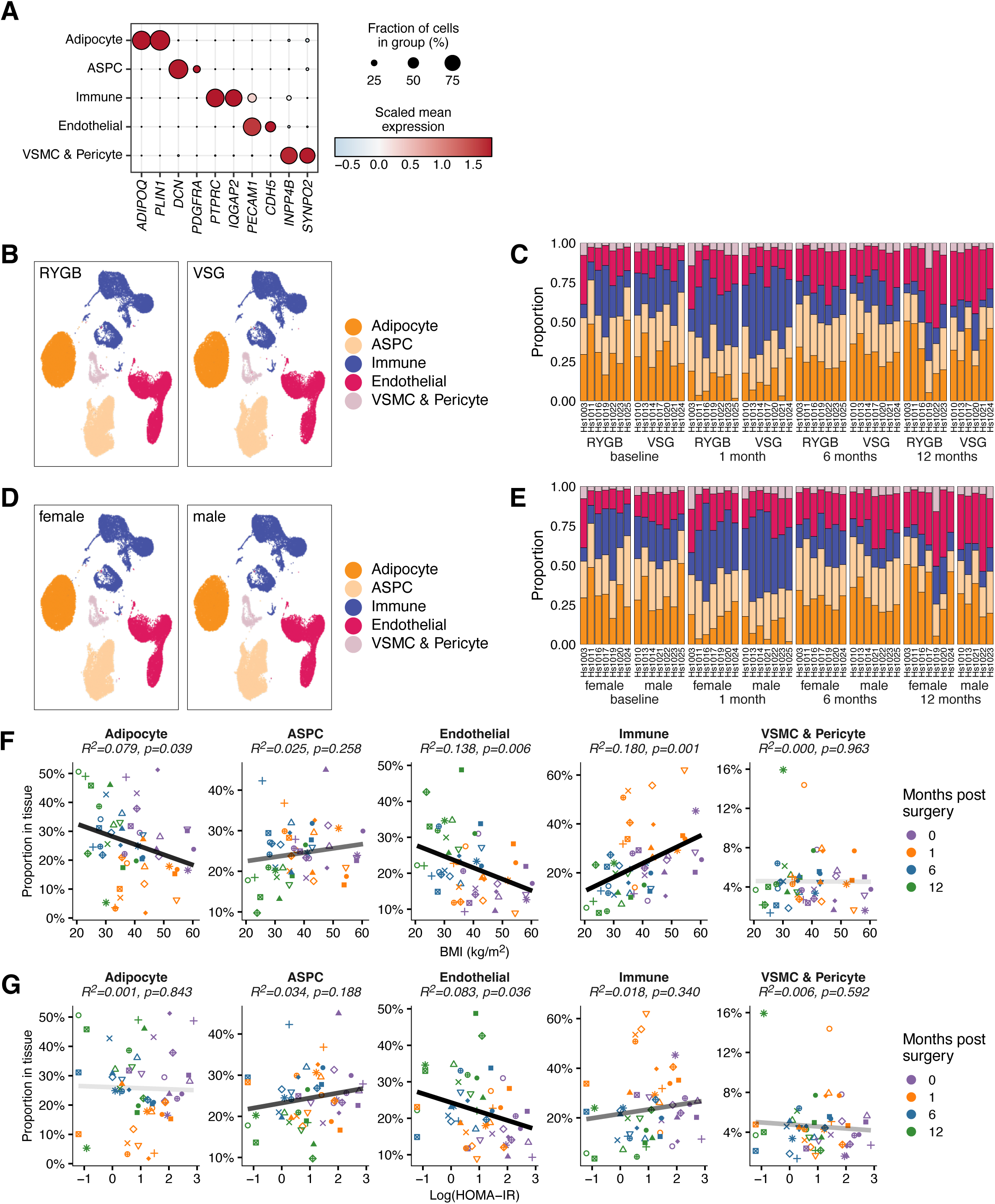
Major cell types in SAT, related to main Figure 1. (A) Expression of broad cell type markers in human SAT. (B) UMAP projections of major cell types in SAT, separated by surgery type. Each plot is normalized to an equal cell count (n = 54,048). (C) Stacked bar plot depicting cell-type composition of each adipose tissue sample, grouped by surgery type and timepoint. Colors reflect cell types in (B). (D) UMAP projections of adipose tissue major cell types, separated by sex. Each plot is normalized to an equal cell count (n = 56,405). (E) Stacked bar plot depicting cell-type composition of each adipose tissue sample, grouped by sex and timepoint. Colors reflect cell types in (D). (F) The relationship between the proportion of each major cell type and the corresponding donor’s BMI at the time of sampling. Individual participants are differentiated by point shape, and different timepoints are indicated by color. (G) The relationship between the proportion of major cell types and the corresponding donor’s HOMA-IR score at the time of sampling. Individual participants are differentiated by point shape, and different timepoints are indicated by color.

**Figure S3.**
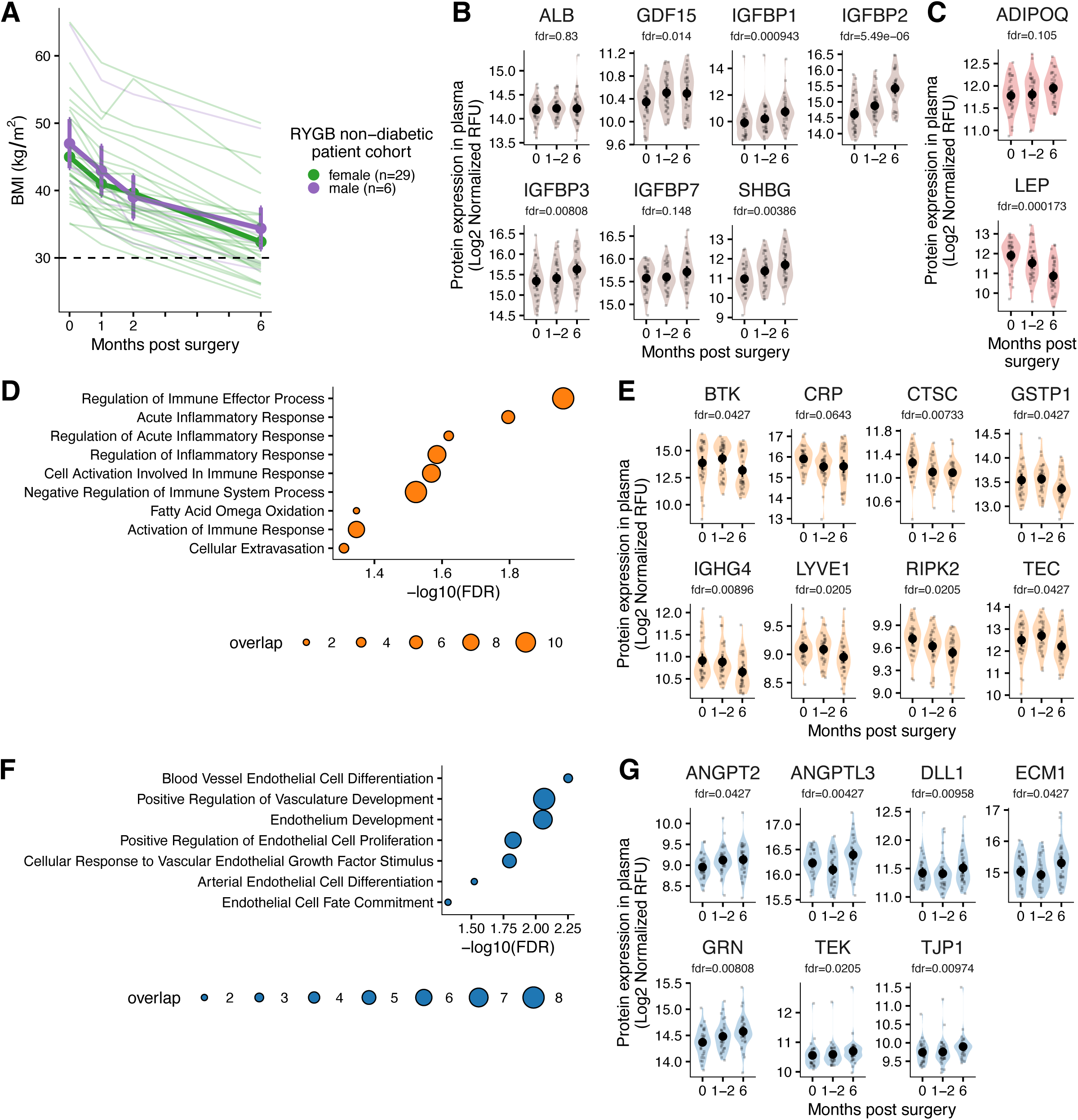
Longitudinal plasma proteomic profiling of bariatric surgery patients, related to main Figures 2 and 3. (A) BMI of SomaScan plasma proteomic study participants following surgery, grouped by sex. The analysis for this manuscript was restricted to the subset of participants who were non-diabetic and had data available across all three study timepoints (n=35). Thin colored lines represent individual participants; thick lines and data points indicate aggregated results across participants. (B) Levels of plasma proteins that reflect participants’ insulin sensitivity after bariatric surgery. Repeated measures ANOVA was used to evaluate statistical significance. Data distribution shown via violin plots (mean ± 95% bootstrapped CI). (C) Levels of plasma adiponectin (ADIPOQ) and leptin (LEP) after bariatric surgery. Repeated measures ANOVA was used to evaluate statistical significance. Data distribution shown via violin plots (mean ± 95% bootstrapped CI). (D) Downregulated inflammatory pathway following bariatric surgery, derived from the list of all significantly downregulated plasma proteins. (E) Levels of plasma proteins associated with inflammatory pathways following bariatric surgery. Repeated measures ANOVA was used to evaluate statistical significance. Data distribution shown via violin plots (mean ± 95% bootstrapped CI). (F) Upregulated vascular development pathways following bariatric surgery, derived from the list of all significantly upregulated plasma proteins. (G) Levels of plasma proteins associated with vascular development following bariatric surgery. Repeated measures ANOVA was used to evaluate statistical significance. Data distribution shown via violin plots (mean ± 95% bootstrapped CI).

**Figure S4.**
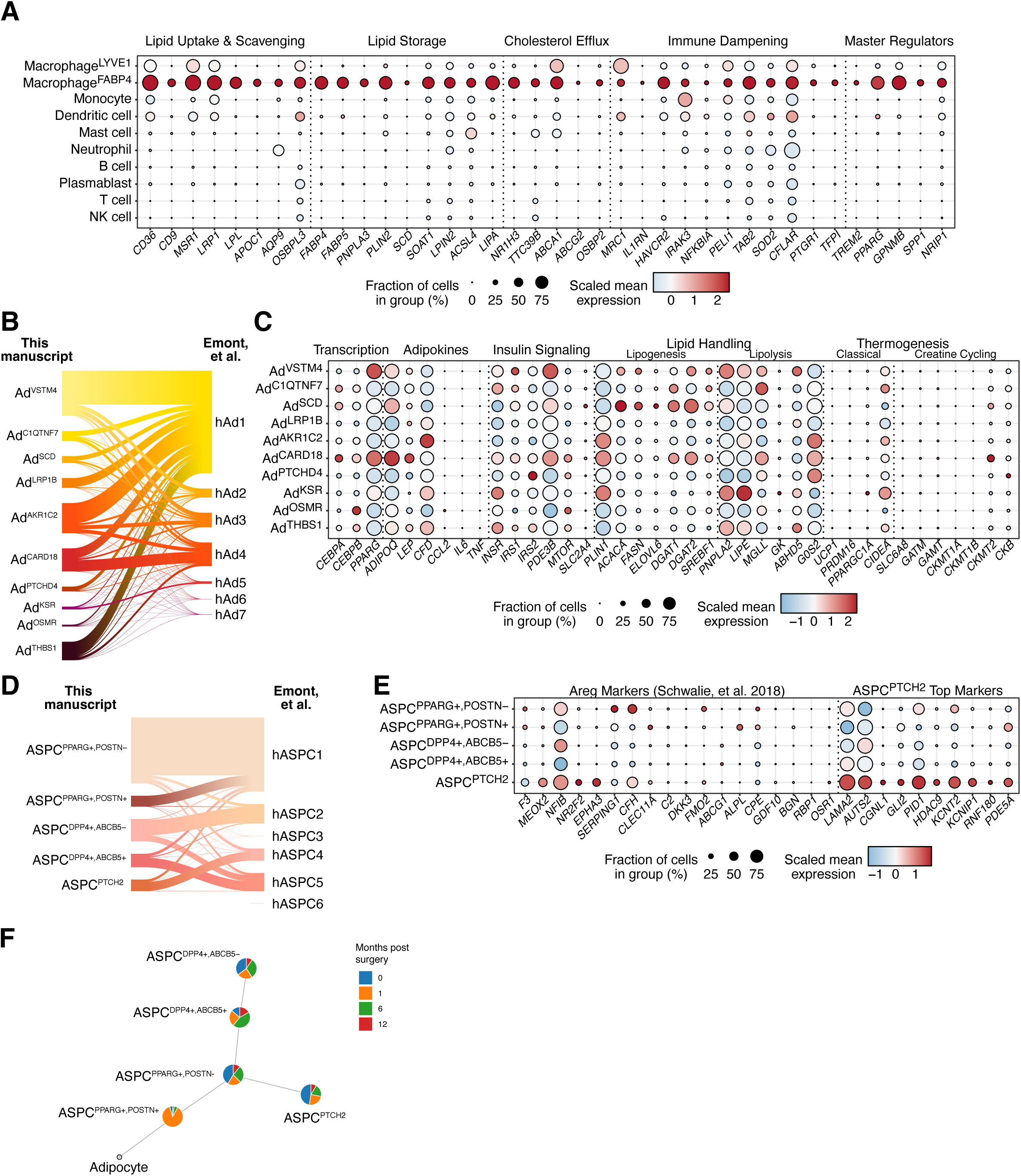
Adipocyte and ASPC subtypes, related to main Figures 4 and 5. (A) Expression of lipid handling genes among Macrophage^FABP4^ markers in immune cells. (B) Riverplot depicting the adipocyte subtype labels used in this manuscript, mapped to the corresponding cell types identified in human SAT adipocyte nuclei from Emont, et al. (C) Expression of genes associated with “transcription”, “adipokine secretion”, “insulin signaling”, “lipid handling”, and “thermogenesis” pathways across adipocyte subtypes. (D) Riverplot depicting the ASPC subtype labels used in this manuscript, mapped to the corresponding cell types identified in human SAT ASPC nuclei data from Emont, et al. (E) Expression of Areg marker genes reported by Schwalie, et al. and selected ASPC^PTCH2^ markers across ASPC subtypes. (F) Lineage structure of ASPC subtypes progressing to adipocytes. Each node contains a pie chart that illustrates the relative abundance of each ASPC subtypes at each timepoint, specifically depicting the percentage of each subtype within the total ASPC cell population.

**Figure S5.**
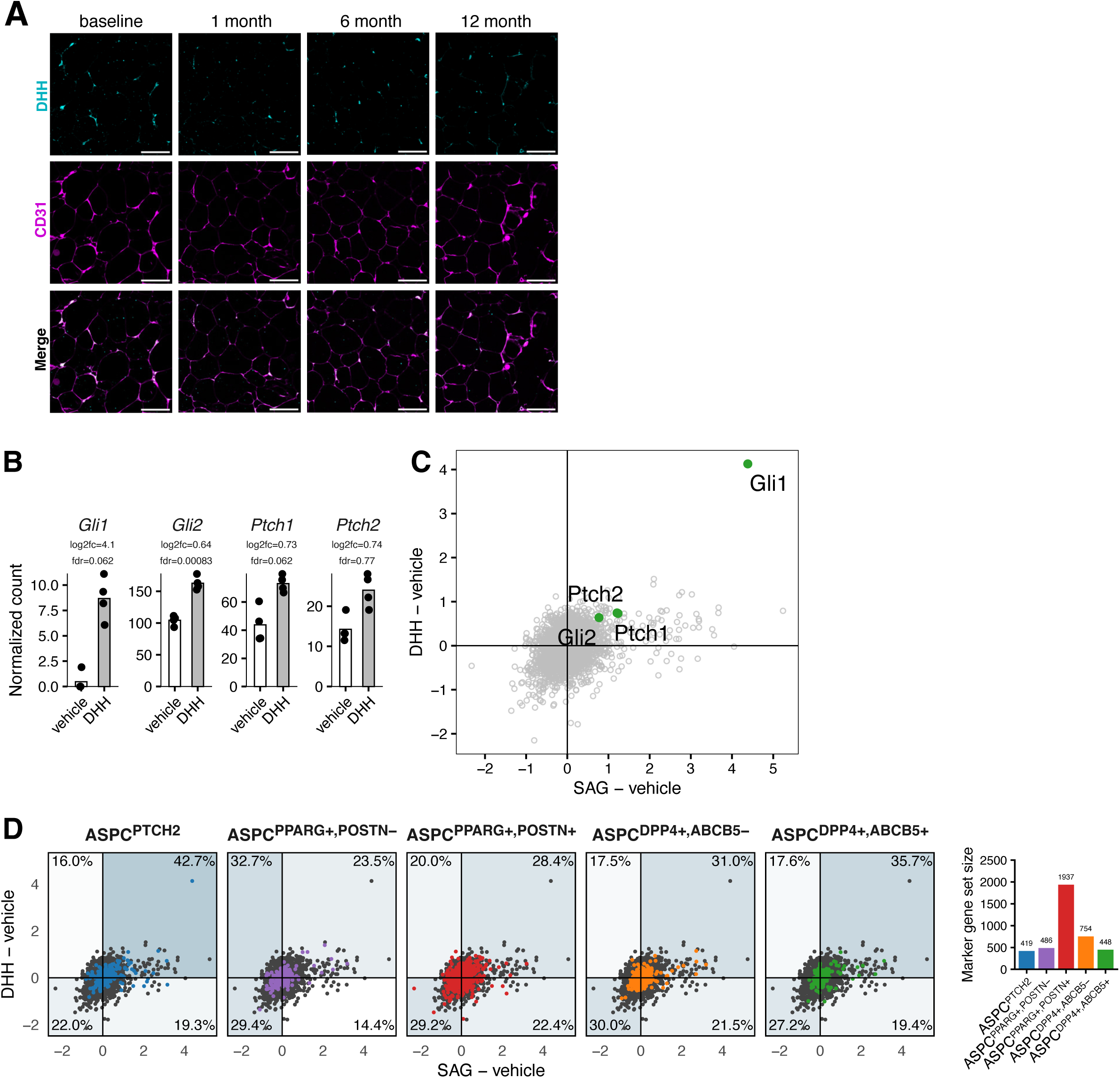
Hedgehog signaling defines ASPC^PTCH2^ identity, related to main Figure 6. (A) Immunofluorescence staining for DHH and pan-endothelial marker CD31 in SAT following bariatric surgery. Scale bar, 100 μm. (B) Expression of hedgehog pathway genes in primary mouse SVF treated with 1 μg/mL of recombinant DHH or vehicle for 24 hours. (C) Scatter plot comparing gene expression changes (log2 fold changes) between SAG-and vehicle-treated cells with those between DHH- and vehicle-treated cells. This plot serves to identify common upregulated signals across the two experiments, with specific highlighting of genes associated with the hedgehog pathway. (D) Scatter plot comparing gene expression changes (log2 fold changes) between SAG-treated and vehicle-treated cells with those between DHH-treated and vehicle-treated cells. Marker genes for each ASPC subtype were highlighted in each plot. The background color in each quadrant of the plot reflects the proportion of subtype marker genes found within that respective quadrant.

**Figure S6.**
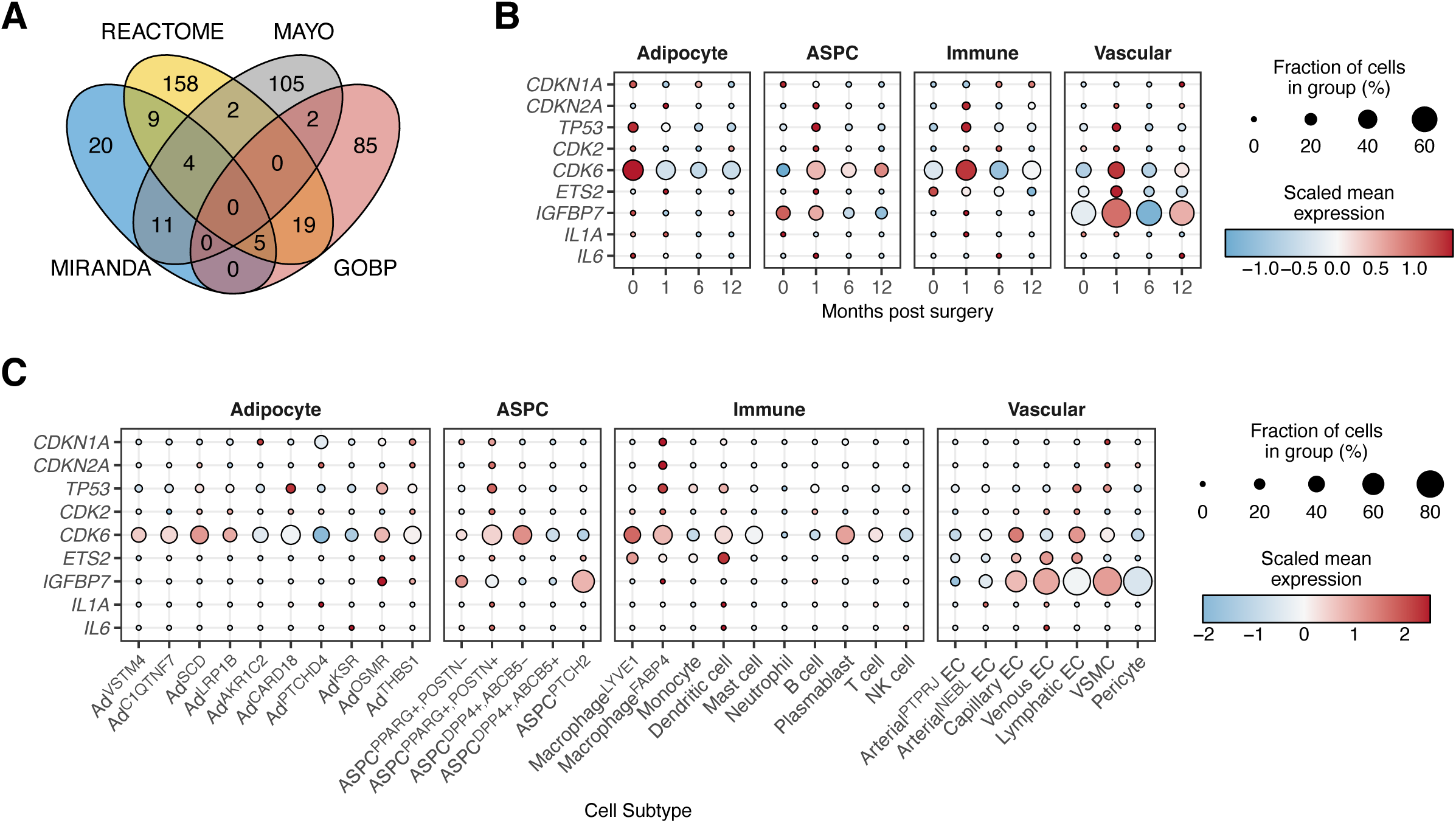
Turnover of senescent adipocytes accompanies metabolic improvement. (A) Venn diagram showing the overlap of genes derived from four distinct senescence gene sets; see text for details. (B) Expression of core senescence-associated genes in major cell types across timepoints. (C) Expression of core senescence-associated genes across cellular subtypes.

## KEY RESOURCES TABLE

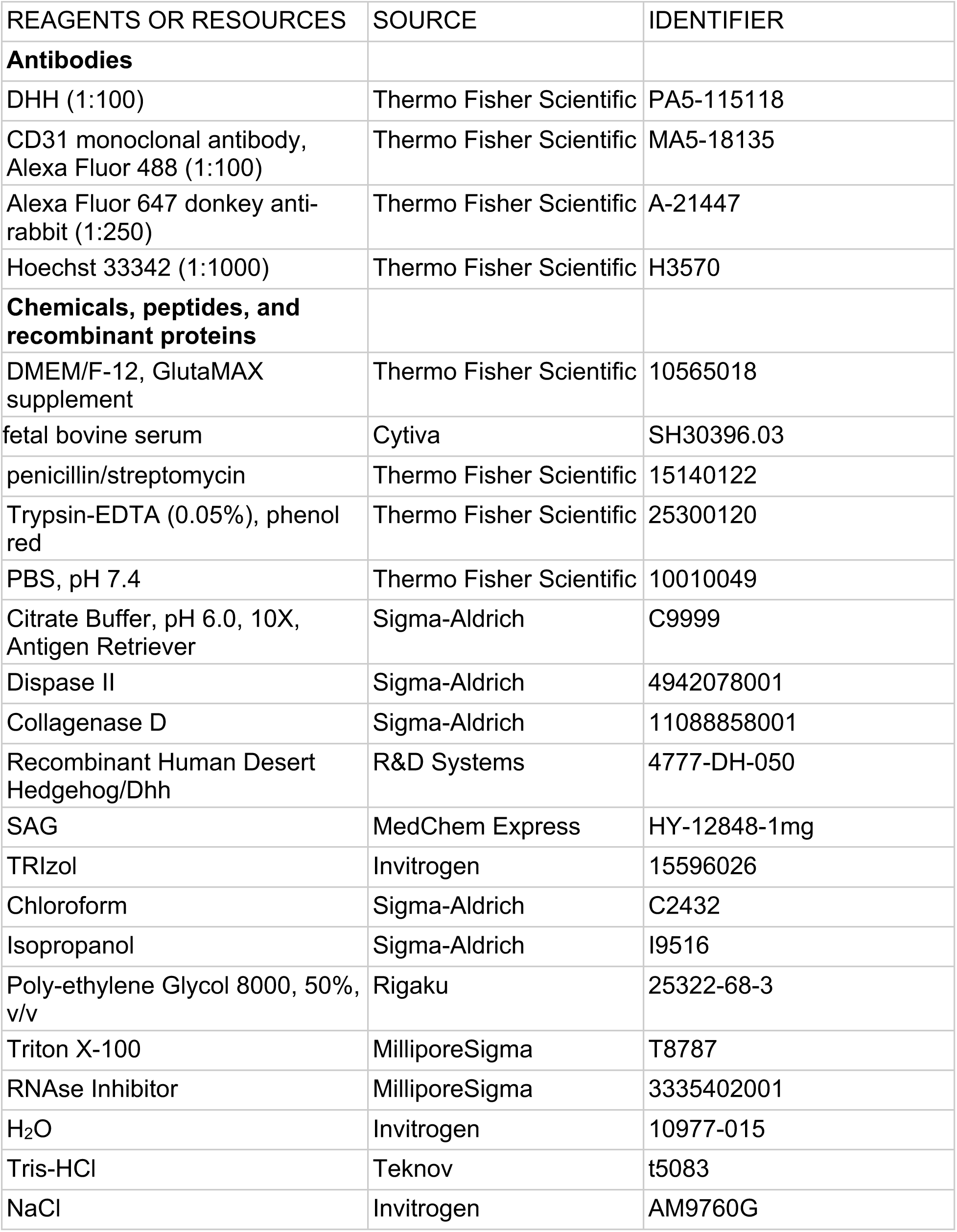
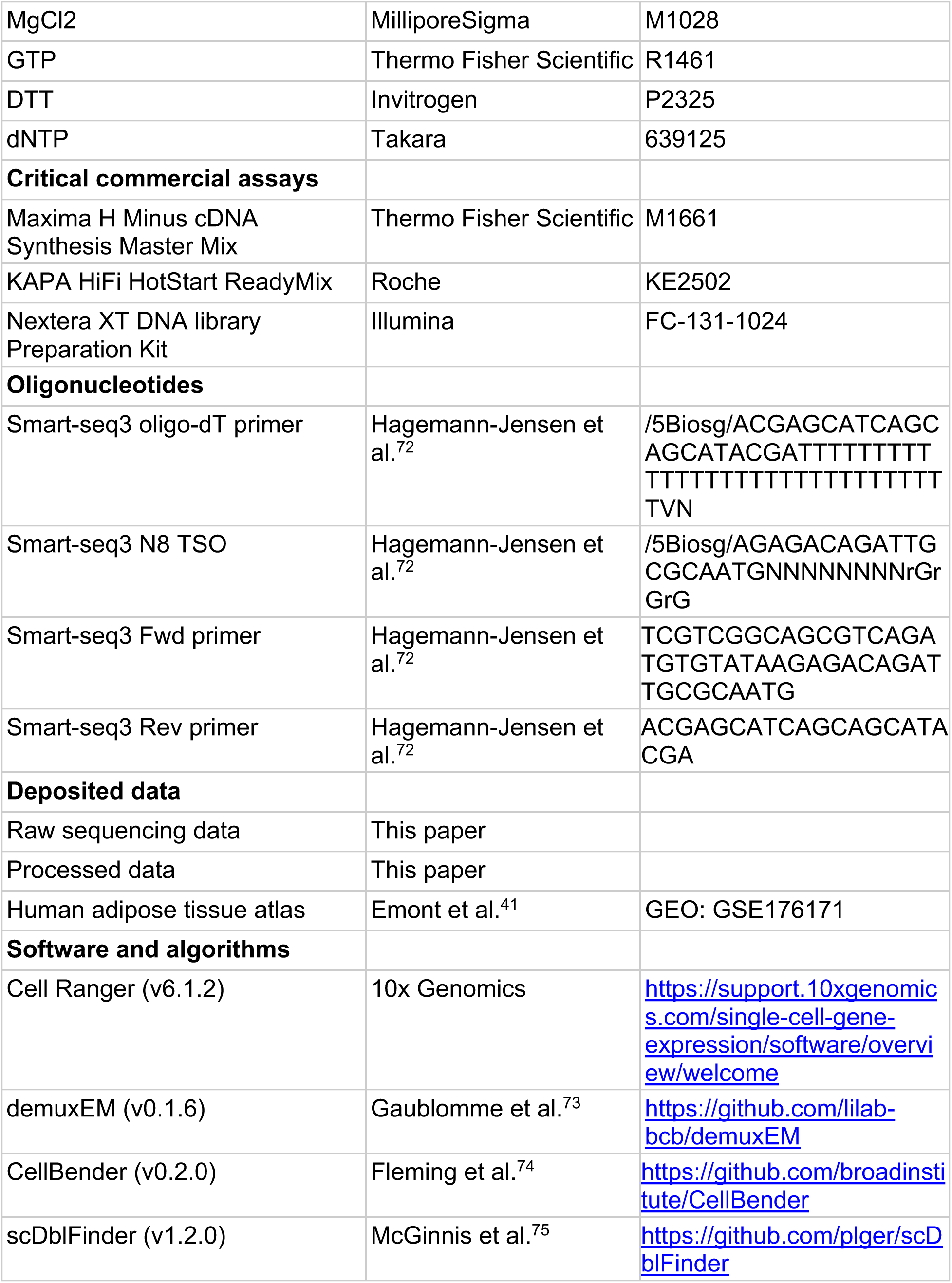
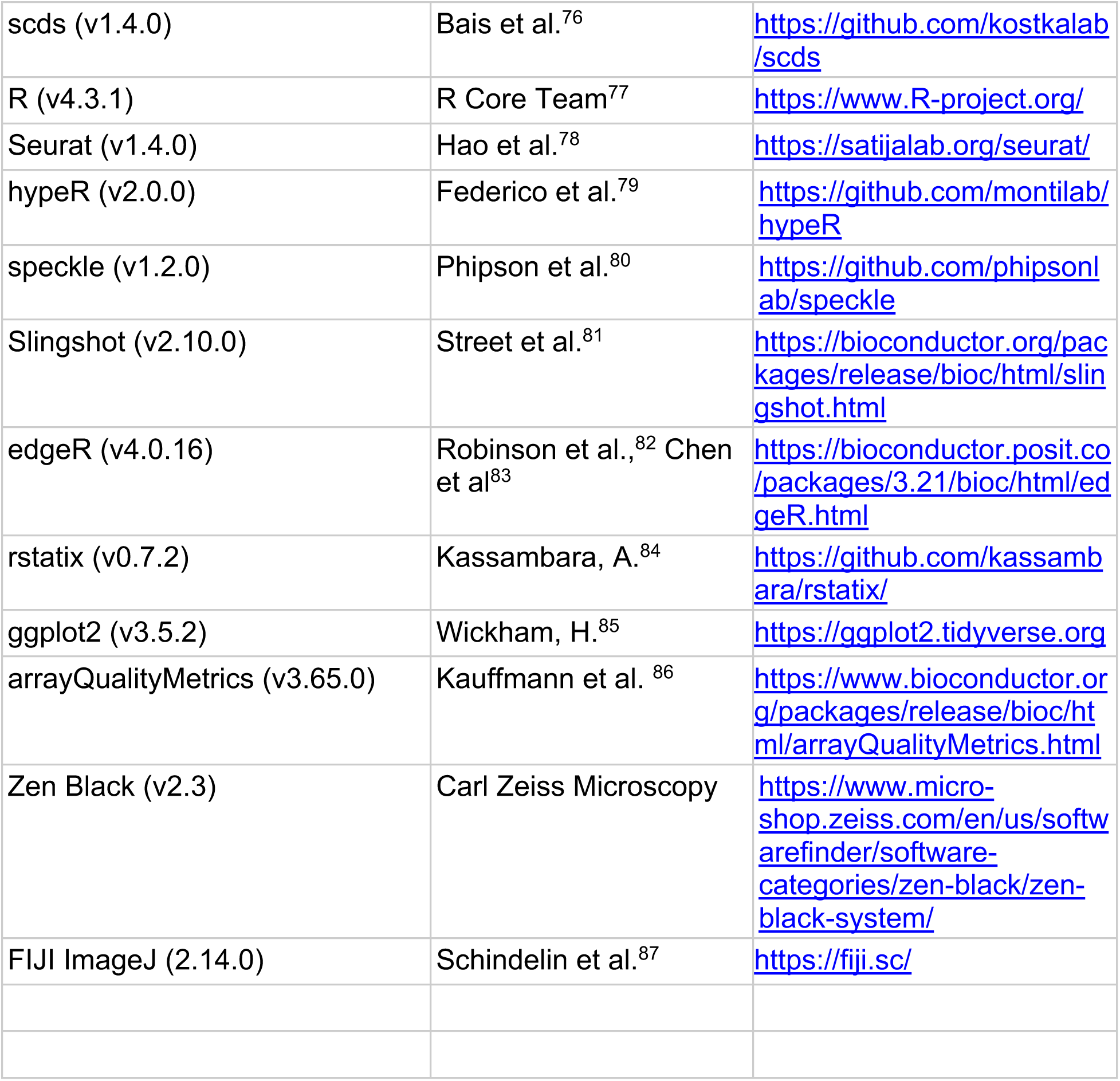

## RESOURCE AVAILABILITY

### Lead contact

Requests for further information should be directed to and will be fulfilled by the lead contact, Evan Rosen.

### Data and code availability

All raw and processed sequencing data generated as part of this study will be deposited at GEO and dbGap and made publicly available at the date of publication. Accession number will be listed in the key resources table. snRNA-seq data will also be available for viewing at the Broad Single Cell Portal (https://singlecell.broadinstitute.org/single_cell/study/SCP3587).

## EXPERIMENTAL MODEL AND SUBJECT DETAILS

### Subject enrollment and sample collection

Subcutaneous adipose tissue was obtained under the University of Pittsburgh Medical Center STUDY 19010309. Patients included were adults aged 21-60 undergoing bariatric surgery. Exclusion criteria were diagnosis of type 1 or type 2 diabetes, pregnancy, alcohol or drug addiction, bleeding/clotting abnormalities, or inflammatory abdominal disease. All patients provided written informed consent prior to surgery.

Baseline samples were collected during bariatric surgery, and subsequent samples were obtained via a "small incisional tissue collection approach" during follow-up visits. The periumbilical skin area was anesthetized and a 6-8 mm incision was made to allow biopsy of a small piece of adipose tissue approximately 0.5-1 cc. The dermal edges were closed with sutures and a dressing was applied. The collected subcutaneous adipose tissue, approximately 0.5-1 cc (0.5 - 1 gram), was sectioned into pieces, approximately 0.25 - 0.5cc (250–500 mg), placed into cryovials, and immediately flash-frozen for subsequent processing.

### Plasma profiling subject enrollment and sample collection

Fasting plasma samples were collected from patients obtaining bariatric surgery from Massachusetts General Hospital. The cohort included men and women of age >18 years who underwent Roux-en-Y gastric bypass at the time of bariatric surgery. Plasma was collected at baseline, 1-2 months and 6 months post-operatively. Patients with pre-existing cardiac disease, chronic kidney disease and uncontrolled hypertension were excluded. For this manuscript, we included results exclusively from patients who did not have diabetes and had plasma measurements recorded at all three timepoints.

### Primary adipose progenitor culture

Freshly harvested mouse inguinal adipose tissues were minced and digested in a solution containing collagenase D (1.5 U/mL) and dispase II (2.4 U/mL), supplemented with 10 mM CaCl₂, for 15–20 minutes in a 37 °C water bath with agitation. Following digestion, wash medium (DMEM-GlutaMAX supplemented with 10% fetal bovine serum and 1% penicillin–streptomycin) was added. The digested tissue was sequentially filtered through 100 μm and 40 μm strainers, and the stromal vascular fraction (SVF) was collected and plated in growth medium (DMEM-GlutaMAX supplemented with 15% FBS and 1% penicillin–streptomycin). After 5 days of culture, adherent preadipocytes were trypsinized and replated at a density of 100,000 cells per well in collagen-coated 12-well plates. Twenty-four hours after plating, cells were treated with either the Hedgehog agonist SAG (200 nM), recombinant DHH (1 μg/mL), or the respective solvent control for 24 hours prior to RNA harvest.

### Mouse VSG

C57BL/6J DIO male mice, 12-13 weeks of age, were purchased from Jackson Labs and maintained on 60% HFD (Research Diets Inc, Cat#D12492) for an additional 2-4 weeks prior to surgery. Body composition was measured using EchoMRI (Echo Medical Systems) before surgery and at the end of study. Sham and VSG surgeries were performed as described previously^34,35^. Briefly, mice were fasted overnight prior to surgery. Animals were anesthetized using isoflurane, and a small laparotomy incision was made in the abdominal wall. The lateral 80% of the stomach along the greater curvature was excised in VSG animals by using an ETS 35-mm staple gun (Ethicon Endo-Surgery). Sham surgery was performed by the application of gentle pressure on the stomach with blunt forceps for 15 seconds. All mice received one dose of Buprinex (0.1 mg/kg) and Carprofen (5 mg/kg) immediately after surgery. All mice received Carprofen (5 mg/kg) for 3 days after surgery. Animals were placed on DietGel Boost (ClearH_2_O; Portland ME) for 3 days after surgery. They were placed back on the pre-operative solid diet (60% HFD) on day 4 post-surgery. Body weight and food intake as well as overall health were monitored daily for the first 7 days after surgery and once weekly until the end of the studies. Semaglutide (0.04 mg/kg) was subcutaneously administered to some Sham-operated mice once daily starting on day 5 post-surgery.

Animals were sacrificed at two timepoints post-surgery, at 10 days for all groups (Timepoint 1), and at 25 days for semaglutide-treated animals and at 35 days for sham and VSG animals (Timepoint 2). All mice were sacrificed on the same calendar days with the order of sacrifice counter-balanced across all the experimental groups.

Because we knew that weight loss would occur faster with Sema, we chose to dose them for 25 days to more carefully match the weight loss between the Sema and VSG-treated mice, their surgeries were performed 10 days later than the others, so that at sacrifice, all animals would be the same age. All mice were fasted for 4 hours prior to tissue collection. Terminal fasting (4 hours) glucose and insulin levels were measured at the end of the study for each cohort.

The presence of contained leaks is assessed at sacrifice in a blinded manner by three experienced surgeons. Each surgeon assigns a score to each particular sleeve and then an average score (out of three values) is calculated. Scores of 1 are assigned when the gastric sleeve is intact and similar in size to that at the time of surgery, with no adhesions present. Scores of 2-3 are assigned for a small, localized leak. Any adhesions are minor. Score 4 is assigned when the sleeve is slightly enlarged but has a single larger leak, with more moderate adhesions to the surrounding organs. Scores of 5 -7 are assigned when there is a large solitary leak to a very large leak with multiple chambers or multiple big leaks. Adhesions range from well-established to very severe. Mice with scores of 4-7 are excluded from the initial analysis primarily due to their pathomorphological changes and inflammatory profile.

## METHOD DETAILS

### Single-nucleus RNA sequencing and data analysis

Flash-frozen human adipose tissue samples (50-300 mg) were homogenized in gentleMACS C Tubes using a gentleMACS Tissue Dissociator with the "mr_adipose_01" program in ice-cold TST buffer, followed by sequential filtration through 40 μm and 20 μm filters. Tissue homogenates were centrifuged at 500g for 5 min at 4°C, washed in nuclei resuspension buffer (NRB), and stained with NucBlue. Individual samples were labeled with unique hashtag antibodies (TotalSeq, 2 μg/mL final concentration) through 45-minute incubation on ice, enabling multiplexing of up to 24 samples per 10X lane. Nuclei were sorted using fluorescence-activated nuclear sorting with gates for single nuclei. Sorted nuclei pools (∼52,000 events targeting ∼40,000 nuclei for loading) were directly loaded onto a 10X Chromium Controller following standard Single Cell 3’ Gene Expression workflow with feature barcoding modifications. Endogenous cDNA and hashtag (HTO) libraries were pooled at 85:15 ratio and sequenced at ≥20,000 reads per nucleus using standard 10X read structure. Raw sequencing reads were processed with Cell Ranger using annotation GENCODE-v44, and sample demultiplexing was performed using demuxEM. Outputs were further processed with CellBender, and doublet scores were calculated using scDblFinder and scds. Cells were removed as doublets if they were both determined to be a doublet using scDblFinder and if they had a scds hybrid score > 1.5.

Count results were further analyzed with R and Seurat. Cells with <800 UMIs and with > 10% of mitochondrial reads were removed from the data object. Genes found in fewer than 2 cells were removed. The data were then normalized using SCTransform and integrated using RPCA in Seurat, integrating by individual so as not to overcorrect across timepoints. To subcluster, cells were subset into broad cell types and reintegrated and clusters were recalculated. Subclusters with high doublet scores as caculated by scDblFinder and scds were removed and subclusters were reintegrated.

The final data set contains 112,991 cells with a median of 3,616 UMIs/cell and a median of 2,003 genes/cell. Reference mapping was performed between the reported dataset and the human subcutaneous dataset from Emont et. al.^41^ using Seurat multimodal reference mapping. Marker genes were calculated using the Seurat Wilcoxon rank sum test with default settings. Pathway analyses were performed using the Bioconductor hypeR package. Statistical assessment of variations in cell type proportions across different timepoints was calculated using the propeller method implemented in the speckle R package. To assess differential gene expression across post-surgical timepoints, single-cell counts were pseudobulked using Seurat’s AggregateExpression function and analyzed via the edgeR Quasi-Likelihood framework. To account for the repeated-measures design, the analysis utilized an additive generalized linear model that tested for timepoint effects while using patient ID as a blocking factor. Raw p-values were adjusted using the Benjamini-Hochberg method. The trajectory of adipocyte development was built by constructing the minimum spanning tree on the Principal Component Analysis (PCA) embeddings of the ASPC subtypes and the adipocytes, using the Bioconductor Slingshot package.

### SomaScan microarray data analysis

The SomaScan assay includes 5,034 SOMAmers, of which 4,783 measure human proteins from 4,137 distinct human genes A hybridization array to capture SOMAmers quantitatively determines the protein present by converting the assay signal (relative fluorescence units) into the relative abundance of an analyte. Assays were performed by SomaLogic in collaboration with Novartis.

R/Bioconductor package arrayQualityMetrics was used for microarray technical quality assessment, and data was background corrected and normalized by SomaLogic using spike in controls and calibrator samples. Differentially measured protein levels were calculated using the repeated measured ANOVA method in the rstatix R package. The p-values were adjusted using Benjamini-Hochberg method to control for the false discovery rate (FDR). Proteins were considered significantly changed if they had an FDR-adjusted p value <0.05 and a fold change greater than 20%.

For the analysis in this manuscript, we used a subset of the original dataset, including only profiles from non-diabetic donors. Furthermore, we restricted the data to only those profiles with complete measurements across all three time points (n=35).

### Immunofluorescence and blood vessel quantification

Flash-frozen adipose tissue biopsies were partially thawed from -80°C to -20°C for 24 hours, then fixated in 10% formalin overnight at 4°C under constant rotation. Fixation was followed by paraffin embedding, sectioning, and staining with hematoxylin and eosin (H&E) to assess section quality. Unstained sections for immunofluorescence were deparaffinized and rehydrated by heating to 55°C and sequential washing in xylene, 100%, 95%, 70%, 50% ethanol and deionized water. Antigen retrieval was performed in sub-boiling 10 mM sodium citrate buffer for 10 min. Sections were washed with 1% donkey serum in PBS with 0.4% Triton X-100 (PBS-T) twice, and blocked with 5% donkey serum in PBS-T for 30 minutes. Sections were then incubated with the primary antibodies diluted in 1% donkey serum PBS-T at room temperature for 1 h and then at 4°C overnight. Primary antibodies and dilutions used for labeling were: Alexa Fluor 488 mouse anti-CD31 (1:100) and rabbit anti-DHH (1:100). Samples were subsequently incubated with secondary antibody Alexa Fluor 647 donkey anti-rabbit (1:250) and Hoechst 33342 (1:1000) diluted in PBS-T at room temperature for 1 hour. Images were obtained using a Zeiss LSM 880 laser scanning confocal system running Zen Black software. Images were analyzed using ImageJ Fiji software. To quantify DHH^+^ blood vessels, tissue sections were fluorescently immunolabeled for CD31 and DHH as above. Five equally powered fields per sample were evaluated for quantification. DHH^+^CD31^+^ double-positive vessel segments were quantified as a percentage of total CD31^+^ vessel segments.

### RNA isolation for bulk RNA-sequencing

Cells in culture wells were collected with TRIzol reagent. The cell-TRIzol mixtures were transferred to collection tubes and homogenized with a Tissuelyser II (QIAGEN). Chloroform was added in a 1:5 ratio by volume and phase separation was performed. The RNA-containing layer was transferred to new tubes, mixed with an equal volume of 100% isopropanol and incubated overnight at −80 °C for precipitation. Precipitated RNA was with 80% ethanol and resuspended in nuclease-free water. RNA concentration and purity were determined using a NanoDrop 2000 (Thermo Scientific). RNA was sent for bulk RNA-sequencing by the Boston Area Diabetes Endocrinology Research Center (BADERC) Functional Genomics Core.

### Bulk RNA-sequencing and data analysis

Bulk RNA-sequencing data was generated following the Smart-seq3 protocol described by Hagemann-Jensen, M.^72^ Samples were reverse transcribed and template switched using the Maxima H Minus cDNA Synthesis Master Mix and the cDNA were subsequently amplified using Kapa HiFi HotStart ReadyMix. Library construction was performed with the Nextera XT DNA Library Preparation Kit. Differential expressions between treatments and controls were evaluated with the Wald test implemented in the DESeq2 R package. Enrichment analysis was performed using the Bioconductor hypeR package.

